# Identification of a secreted protease from *Bacteroides fragilis* that induces intestinal pain and inflammation by cleavage of PAR_2_

**DOI:** 10.1101/2025.01.15.633241

**Authors:** Markus Lakemeyer, Rocco Latorre, Kristyna Blazkova, Dane Jensen, Hannah M. Wood, Nayab Shakil, Scott C. Thomas, Deepak Saxena, Yatendra Mulpuri, David Poolman, Paz Duran de Haro, Laura J. Keller, David E. Reed, Brian L. Schmidt, Alan E. Lomax, Nigel W. Bunnett, Matthew Bogyo

**Author notes:** These authors contributed equally (Markus Lakemeyer, Rocco Latorre, Kristyna Blazkova).

## Abstract

Protease-activated receptor 2 (PAR_2_) is a central regulator of intestinal barrier function, inflammation and pain. Upregulated intestinal proteolysis and PAR_2_-signaling are implicated in inflammatory bowel diseases (IBDs) and irritable bowel syndrome (IBS). To identify potential bacterial regulators of PAR_2_ activity, we developed a functional assay for PAR_2_ processing and used it to screen conditioned media from a library of diverse gut commensal microbes. We found that multiple bacteria secrete proteases that cleave host PAR_2_. Using chemoproteomic profiling with a covalent irreversible inhibitor, we identified a previously uncharacterized *Bacteroides fragilis* serine protease Bfp1, and showed that it cleaves and activates PAR_2_ in multicellular and murine models. PAR_2_ cleavage by Bfp1 disrupts the intestinal barrier, sensitizes nociceptors, and triggers colonic inflammation and abdominal pain. Collectively, our findings uncover Bfp1-mediated PAR_2_-processing as a new axis of host-commensal-interaction in the gut that has the potential to be targeted for therapeutic intervention in IBD or IBS.

## INTRODUCTION

An imbalance of the gut microbiota, known as dysbiosis, has been linked to gastrointestinal diseases such as colorectal cancer,^1^ inflammatory bowel disease (IBD)^2^ and irritable bowel syndrome (IBS)^3^, as well as seemingly unrelated conditions including Parkinson’s disease^4^ and response to chemotherapy.^5^ Despite the clinical relevance of these observations, the underlying mechanisms that define specific bacterial contributions to pathogenesis remain largely unclear. While individual bacterial metabolites have been shown to be regulators of bacterial-host signaling,^6,7^ the role of secreted proteins, especially bacterial enzymes, has been mostly overlooked.^8^ Intriguingly, the gut is the organ most exposed to proteases, both from endogenous and exogenous sources.^9^ These proteases not only mediate the digestion of dietary proteins but also regulate inflammation, pain, cell migration, apoptosis and intestinal permeability.^10^ Excessive proteolysis in the gut caused by host or microbial proteases has been identified as a major contributor to IBD^2,11^ and IBS.^12^ Moreover, proteases play pivotal roles in microbial homeostasis and mediate microbe-microbe interactions.^8,9^ Therefore, identifying and modulating bacterial protease activities are promising strategies for the development of therapies for microbiome-related diseases.^13,14^

Protease-activated receptor 2 (PAR_2_) is a central regulator of epithelial barrier function and a key modulator of intestinal inflammation and pain. PAR_2_ is a G-protein coupled receptor (GPCR) that mediates a number of downstream signaling pathways. In contrast to most GPCRs, PAR_2_ is not activated by small molecule binding. Rather, proteolysis within the extracellular N-terminal domain (NTD) reveals a neo-N-terminus that acts as a tethered ligand for the cleaved receptor (**Figure 1A**) or induces conformational changes that activate the cleaved receptor.^15^ Excessive proteolysis in the gut and consequent PAR_2_-signaling have been linked to IBD and IBS severity and intestinal pain.^11,12^ While the modulation of PAR_2_ signaling is understood as a virulence strategy for pathogens such as *Enterococcus faecalis*,^15^ it remains unclear which extracellular proteases from co-occurring commensal bacteria modulate host PAR_2_ signaling and thereby affect gut physiology and pathology.

**Figure 1.**
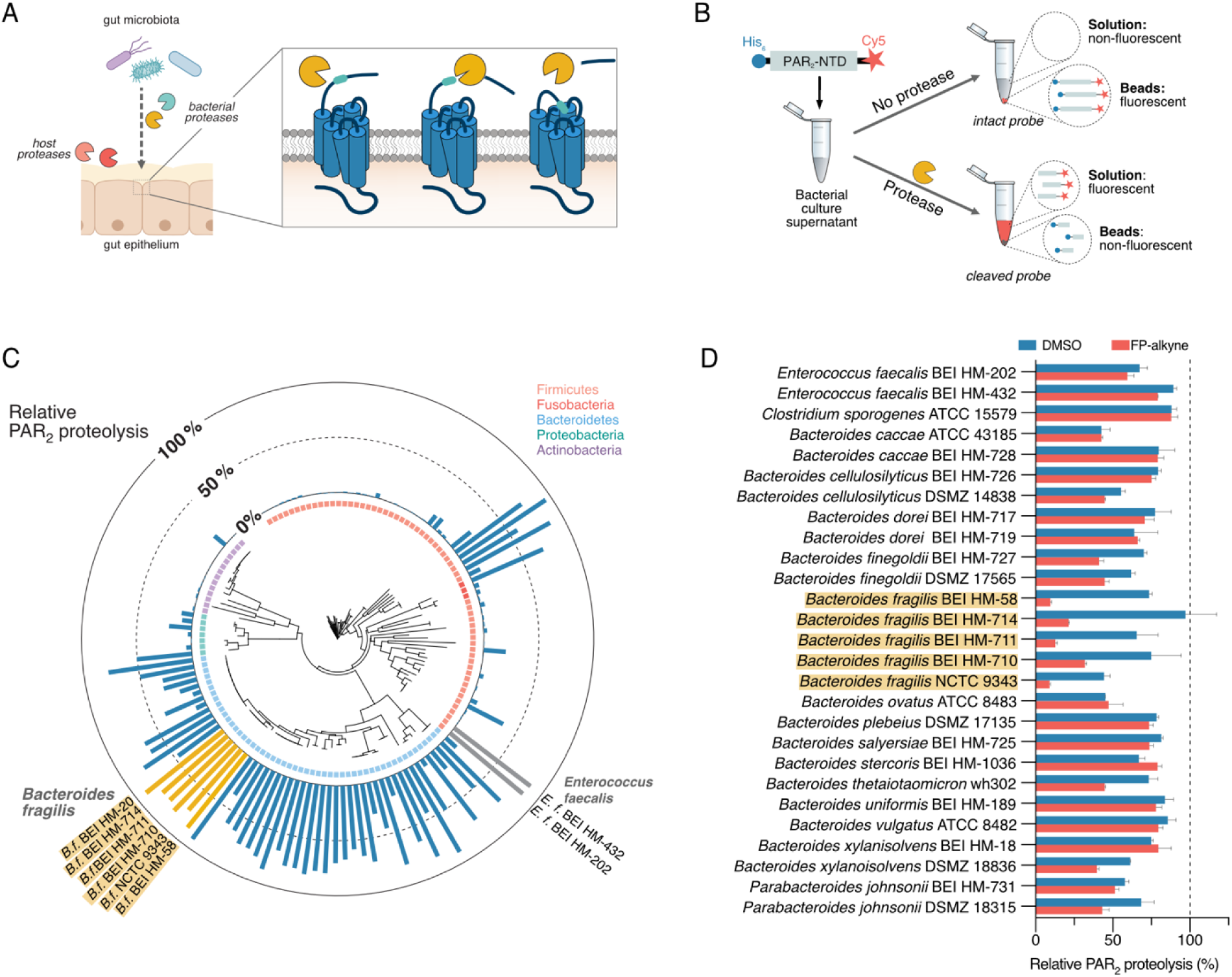
Identification of bacterial proteases that cleave the PAR_2_-NTD. (A) Mechanism of activation of PAR_2_ through host or microbial proteases. (B) Schematic representation of the dual-labeled substrate probe and workflow for the screening of bacterial cultures for PAR_2_-NTD cleaving proteases. After incubation with culture supernatants, intact or cleaved substrate is affinity-enriched on magnetic Ni-NTA beads and fluorescence in solution measured to quantify proteolysis. (C) Results from the PAR_2_-NTD proteolysis screening mapped onto a phylogenetic tree of all bacterial strains tested. Bar graph depicts PAR_2_-NTD proteolysis relative to no-protease control (0% cleavage) and treatment with excess of trypsin (100% cleavage). (D) PAR_2_-processing activity of most active supernatants upon pre-incubation DMSO (control) or serine hydrolase inhibitor fluorophosphonate-alkyne (FP-alkyne, 20 µM). Data are presented as mean ± SEM.

Here, we describe a screening platform for functional profiling of the secretomes of gut bacteria that process human PAR_2_, which identified members of the Bacteroidetes phylum with high PAR_2_-cleaving activity. By applying covalent inhibitors and chemical proteomics, we identified a previously uncharacterized and unique serine protease that we have named Bfp1 from *Bacteroides fragilis*, which efficiently processes PAR_2_, resulting in receptor activation, internalization, and increased transepithelial permeability in multicellular models. Intracolonic administration of *B. fragilis* culture supernatants as well as engraftment of mice with wild-type (WT) *B. fragilis* bacteria resulted in strong PAR_2_-mediated colonic inflammation and nociception that was absent in a Δ*bfp1*-knockout (KO) strain. Colonic administration of Bfp1 *in vitro* led to excitation of sensory nerves and this sensitization was lost in mice lacking PAR_2_ in NaV1.8-expressing nociceptors. Overall, our work has uncovered Bfp1-mediated PAR_2_-processing as a novel mechanism of host-commensal interactions in the gastrointestinal tract that has profound impact on host physiology.

## RESULTS

### Development of an affinity-pulldown platform to profile microbial PAR_2_-proteolysis

To identify bacterial strains that process the human PAR_2_-NTD *via* secreted proteases, we developed a sensitive *in vitro* screening platform for directly testing non-concentrated bacterial culture supernatants in a multiplexed manner. Using expressed protein ligation and bioorthogonal chemistry (**Figure S1** and Supplementary Note), we generated a synthetic dual-labeled reporter substrate representing the 46 amino acid-spanning PAR_2_-NTD with N-terminal hexahistidine (His_6_) affinity tag and C-terminal Cy5 fluorophore (**Figure 1B**). Upon cleavage by microbial proteases, the fluorophore is separated from the affinity tag. When the His_6_-tagged peptides are removed from the solution *via* Ni-NTA-based affinity pulldown, the extent of PAR_2_-NTD cleavage can be quantified by the amount of released fluorescent fragment that remaining in the soluble fraction (**Figure 1B**). Optimization using recombinant trypsin, the canonical PAR_2_ activator, and *Enterococcus faecalis* culture supernatants that contain the known PAR_2_-cleaving protease gelatinase^16^ resulted in a robust and sensitive assay that requires minimal volumes (10 µL) of non-concentrated culture supernatants (**Figure S2**).

We applied our screening platform to a collection of 140 human gut microorganisms, representing 104 species and spanning 5 phyla, that has previously been applied to profile the production of microbial metabolites.^17^ To create comparable datasets, we cultivated all strains in a rich, undefined medium (Mega Medium, MM), which supports the growth of diverse bacteria.^17^ Bacteria were grown in 96-well microtiter plates, harvested by centrifugation and sterile-filtration, and subsequently tested in the PAR_2_-proteolysis assay. As an outcome of this global screening, we identified culture supernatants from 45 bacterial strains that processed more than 50% of the PAR_2_-substrate in our assay (**Figure 1C, Table S1**). While it has previously only been hypothesized that commensals might interact with the human host *via* PAR-modulation, these results highlight that a remarkable number of bacterial strains secrete potential PAR_2_-processing proteases. In addition to select Firmicutes species, such as two *Enterococcus faecalis* strains, we observed very strong proteolytic activities within the Bacteroidetes phylum. In particular, five *Bacteroides fragilis* strains showed consistently high PAR_2_-processing, ranging from 78% – 86% of total substrate. For the validation of their proteolytic activity, all “hit” strains were individually cultivated on a large scale, harvested at early stationary phase (to exclude release of intracellular proteases through cell lysis) and subjected to repeated testing with a shortened proteolysis time (to select for most active supernatants). Out of the 45 initial hits, 27 passed these more stringent selection criteria (**Figure 1D** and **S3**, see Supporting Information). It is noteworthy that apart from the two *E. faecalis* strains, *Clostridium sporogenes* ATCC 15579 remained the only non-Bacteroidetes PAR_2_-processor identified.

### Identification of uncharacterized *B. fragilis* serine proteases by chemical proteomics

Proteases can be classified based on their active sites into serine-, cysteine, metallo-, aspartic-/glutamic-, asparagine- and threonine-proteases. Previous work has linked excessive serine protease activities with intestinal diseases such as IBD and IBS.^8,18,19^ Multiple serine proteases from the host or exogenous sources are known to cleave and activate PAR_2_.^20^ As serine proteases are additionally predicted to be the largest group of proteases to be secreted by bacteria,^21^ and can be efficiently inhibited by broad-spectrum inhibitors, we focused our protein identification efforts specifically on serine proteases.

Bacterial culture supernatants were pre-treated with a covalent, broad-spectrum serine hydrolase probe, fluorophosphonate (FP)-alkyne (**Figure 2A**)^22^, prior to the PAR_2_-proteolysis assay (**Figure 1D**). FP-alkyne partially inhibited PAR_2_ cleavage by select strains, including *Bacteroides finegoldii* BEI HM-727 and *Bacteroides thetaiotaomicron* wh302. The strongest inhibitory effect was observed for five *Bacteroides fragilis* strains, resulting in 58 – 87 % inhibition compared to vehicle (DMSO) -treated samples (**Figure 1D**). We performed activity-based protein profiling (ABPP, schematic workflow in **Figure 2B**)^23^ on culture supernatants of the reference strain *B. fragilis* NCTC 9343 to identify the PAR_2_- processing protease(s). We first optimized labeling conditions *via* gel-based ABPP using a fluorescent FP-tetramethylrhodamine (FP-TMR) probe (**Figure S4**), prior to quantitative ABPP using an FP-biotin probe, streptavidin-based affinity-enrichment of labeled proteins and LC-MS/MS-based analysis. This identified 20 proteins, 10 of which are potential peptidases or proteases, that were strongly (>4-fold) and significantly (*p*<0.05) enriched compared to the vehicle (DMSO) control (**Figure 2C, Table S2**). For the unambiguous identification of the PAR_2_-processing protease, we screened a small set of 11 covalent protease inhibitors (**Figure S5A**) and identified the peptide chloromethyl ketone **MLP7** (**Figure 2A**) as an inhibitor of the PAR_2_-processing activity of *B. fragilis* (**Figure S5B**). Pretreatment of culture supernatants with **MLP7** prior to labeling with FP-biotin blocked the protease-labeling by FP-probes in a competitive and concentration-dependent manner (**Figure 2B and S4**). Importantly, the MS-based competitive ABPP experiment resulted in only two protease candidates with the UniProt ^24^ identifiers Q5LDF9 (gene name BF9343_2070) and Q5LIA5 (gene name BF9343_0342) (**Figure 2D, Table S2**). Both enzymes are uncharacterized putative lipoproteins that, according to InterPro analysis^25^, belong to the poorly characterized families of S41-proteases or tail-specific proteases (**Figure 2E**).

**Figure 2.**
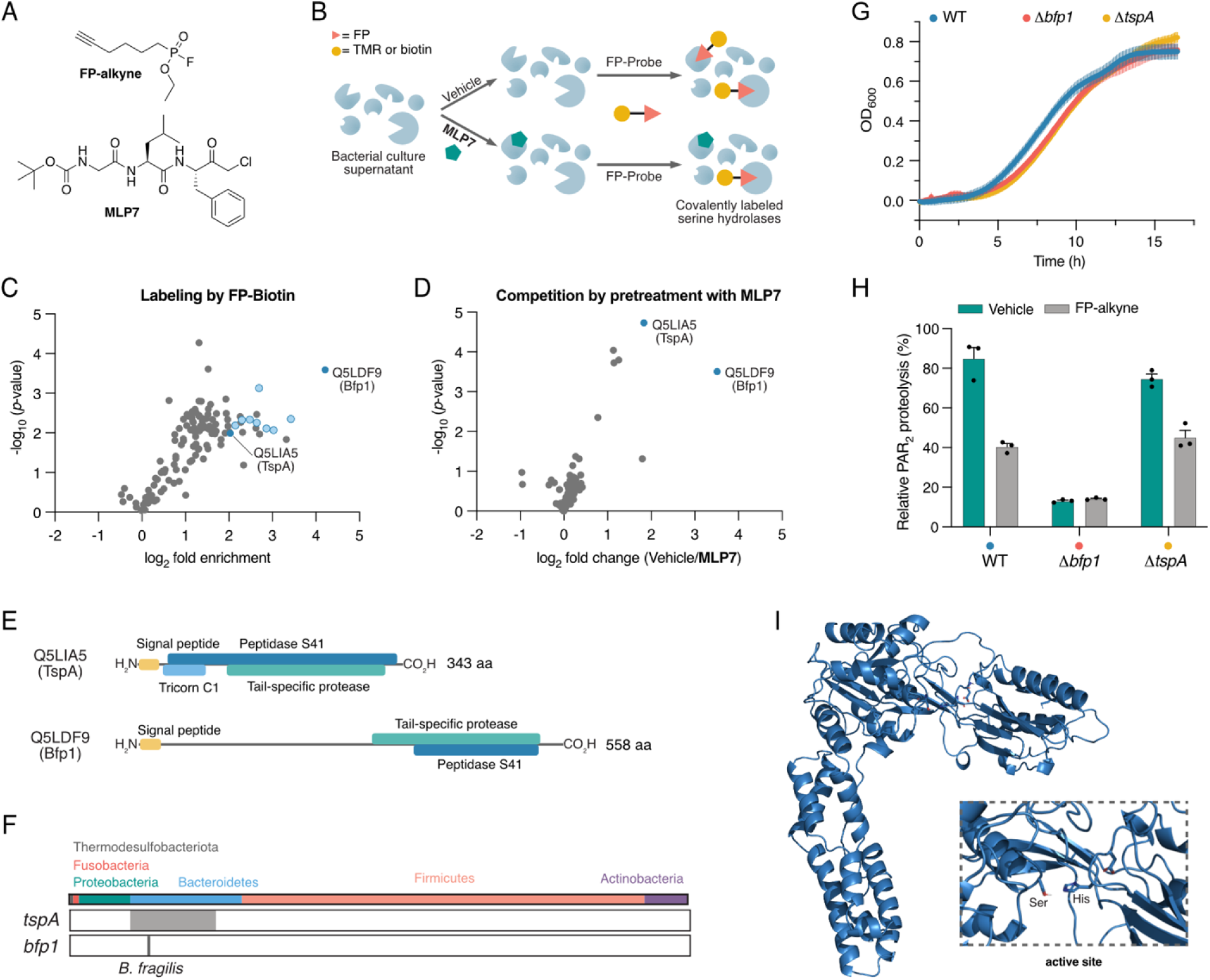
Identification of two uncharacterized *B. fragilis* proteases. (A) Chemical structures of inhibitors **FP-alkyne** and **MLP7**. (B) Schematic representation of competitive ABPP workflow. (C) Volcano plot of FP-biotin protein targets in the secretome of *B. fragilis* NCTC 9343. Significantly enriched peptidase/protease candidates are colored in blue. (D) Volcano plot for competitive ABPP experiment with pretreatment by **MLP7** uncovers two PAR_2_- processing candidates, TspA and Bfp1. (E) InterPro domain analysis of the proteases Q5LIA5 (TspA) and Q5LDF9 (Bfp1). (F) Phylogenetic distribution of *bfp1* and *tspA* based a database of 1,520 reference genomes of culturable bacteria from human gut microbiota. (G) Growth curves for WT *B. fragilis* and the *B. fragilis* Δ*bfp1* and Δ*tspA* protease knockouts. (H) Relative PAR_2_-proteolysis of the WT *B. fragilis* and respective knockout strains confirms that Bfp1 is the responsible PAR_2_-processing protease. (I) Structural prediction of Bfp1 (AlphaFold model) with a typical S41 serine protease domain and an extended helical domain of unknown function.

BLAST analysis against 1,520 reference genomes from cultivated human gut bacteria^26^ resulted in 381 Bacteroidetes spp. with homologs of Q5LIA5 (**Figure 2F, Table S3**). In contrast, Q5LDF9 was only identified in the genomes of the 31 unambiguously annotated *B. fragilis* strains in the database (**Figure 2F, Table S4**). We therefore propose to name the conserved Q5LIA5 protease tail-<u>s</u>pecific <u>p</u>rotease <u>A</u> (**TspA**) and the Q5LDF9 enzyme *<u>B</u>acteroides fragilis* serine <u>p</u>rotease <u>1</u> (**Bfp1**). The unique distribution of Bfp1 in *B. fragilis* and lack of Bfp1-negative *B. fragilis* strains suggests a specific physiological role for this serine protease.

### Genetic deletion mutants uncover Bfp1 to be the PAR_2_-processing protease

To determine which enzyme candidate was responsible for our observed PAR_2_-processing activity, we generated two *B. fragilis* NCTC9343 strains in which either the *tspA* or the *bfp1* gene was genetically deleted by heterologous recombination (Supporting Information).^27^ Both deletion mutants showed only a slightly delayed growth compared to the WT strain, highlighting that these enzymes are not essential for bacterial survival *in vitro* under our conditions (**Figure 2G**). Next, bacterial culture supernatants were subjected to the PAR_2_-proteolysis assay. While knockout of *tspA* did not alter PAR_2_-processing compared to WT *B. fragilis*, the supernatants from the *B. fragilis* Δ*bfp1* strain exhibited only residual PAR_2_-processing activity, which could not be further reduced by pretreatment with FP-alkyne (**Figure 2H**). Consequently, we concluded that Bfp1 is the serine protease that mediates PAR_2_-NTD proteolysis in *Bacteroides fragilis* culture supernatants. The predicted structure of Bfp1^28^ contains a proteolytic core with its active site serine residue and an extended helical domain which is annotated as “domain of unknown function” (**Figure 2I**).^25^

### Bfp1-mediated cleavage of full-length PAR_2_ at the cell-surface

To determine whether bacterial Bfp1 can cleave the extracellular NTD of intact PAR_2_ at the cell surface and stimulate endocytosis of the cleaved receptor, consistent with activation, we incubated *B. fragilis* culture supernatants with HEK293T cells expressing human PAR_2_ with an extracellular HA-epitope and an intracellular fluorescent mApple tag (**Figure 3A**). Cells were exposed to bacterial supernatants, and HA epitope and mApple were localized by immuno-fluorescence microscopy. In unstimulated cells and in cells exposed to Mega Medium only (control), HA and mApple colocalized at the plasma membrane (**Figure 3B**). Trypsin (10 nM, 30 min, positive control) removed the HA-tag and caused redistribution of mApple to endosomes, consistent with PAR_2_ cleavage and activation. A similarly strong phenotype was observed upon treatment with culture supernatants from WT *B. fragilis* and this altered localization was partially restored for *B. fragilis* Δ*bfp1*, highlighting that Bfp1 indeed processes the intact, membrane-bound PAR_2_ (**Figure 3B**).

**Figure 3.**
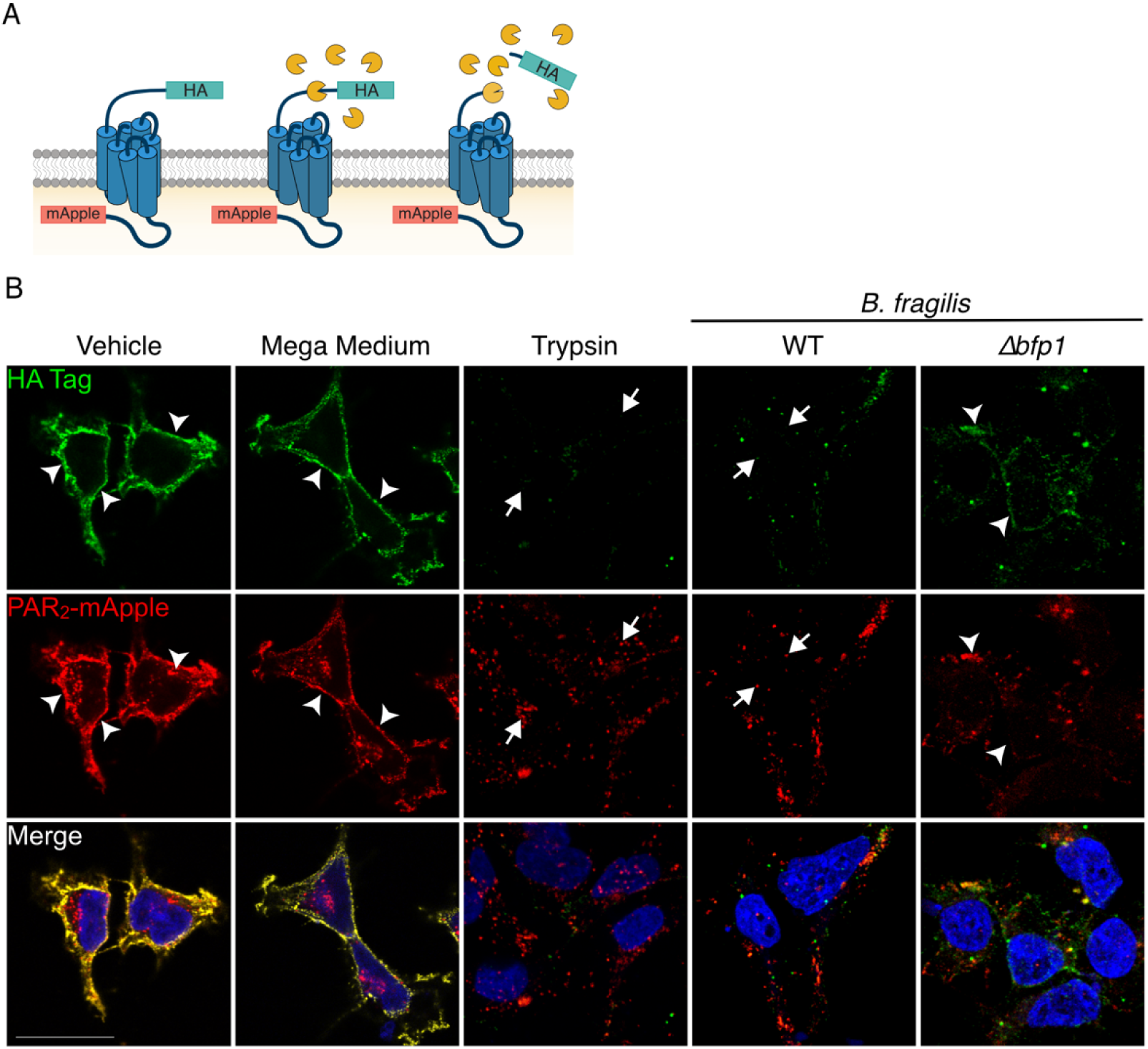
Bfp1 processes PAR_2_ at the cell-surface and induces endocytosis. (A). Validation of N-terminal processing of full-length, membrane-bound PAR_2_ using HEK293T cells expressing PAR_2_ with an N-terminal HA-epitope (green) and C-terminal mApple fluorescent tag (red). (B). Cells were exposed to trypsin (10 nM) or 10-fold concentrated supernatants from WT *B. fragilis* or Δ*bfp1*-knockout cultures, or equally concentrated Mega Medium (MM) and analyzed by immuno-fluorescence microscopy to detect HA and mAppple. Trypsin (positive control) removes Flag and stimulates redistribution of mApple to endosomes (arrowheads), denoting PAR_2_ cleavage, activation and internalization. WT *B. fragilis* supernatant has similar effects. *B. fragilis* Δ*bfp1*-knockout supernatant does not completely remove Flag, denoting slower cleavage of PAR_2_. Scale bar: 20 µm, arrowheads denote surface PAR_2_, arrows show internalized receptor.

### Bfp1 increases epithelial permeability in human intestinal organoids

PAR_2_ is a central modulator of the intestinal epithelial barrier and excessive activation of PAR_2_ has been linked to increased paracellular permeability and “leaky gut” syndrome.^2,9,29^ To assess whether Bfp1 modulates intestinal permeability in a physiologically relevant human model, we subjected organoids from healthy ileal adult tissue to bacterial culture supernatants and analyzed paracellular permeability in organoid-derived monolayers and apical-out organoids (**Figure 4A**). Apical-out organoids facilitate permeability testing as effector molecules that are predicted to interact with the apical side can be added directly to the culture medium, circumventing the need for laborious intra-organoid injection.^30^

**Figure 4.**
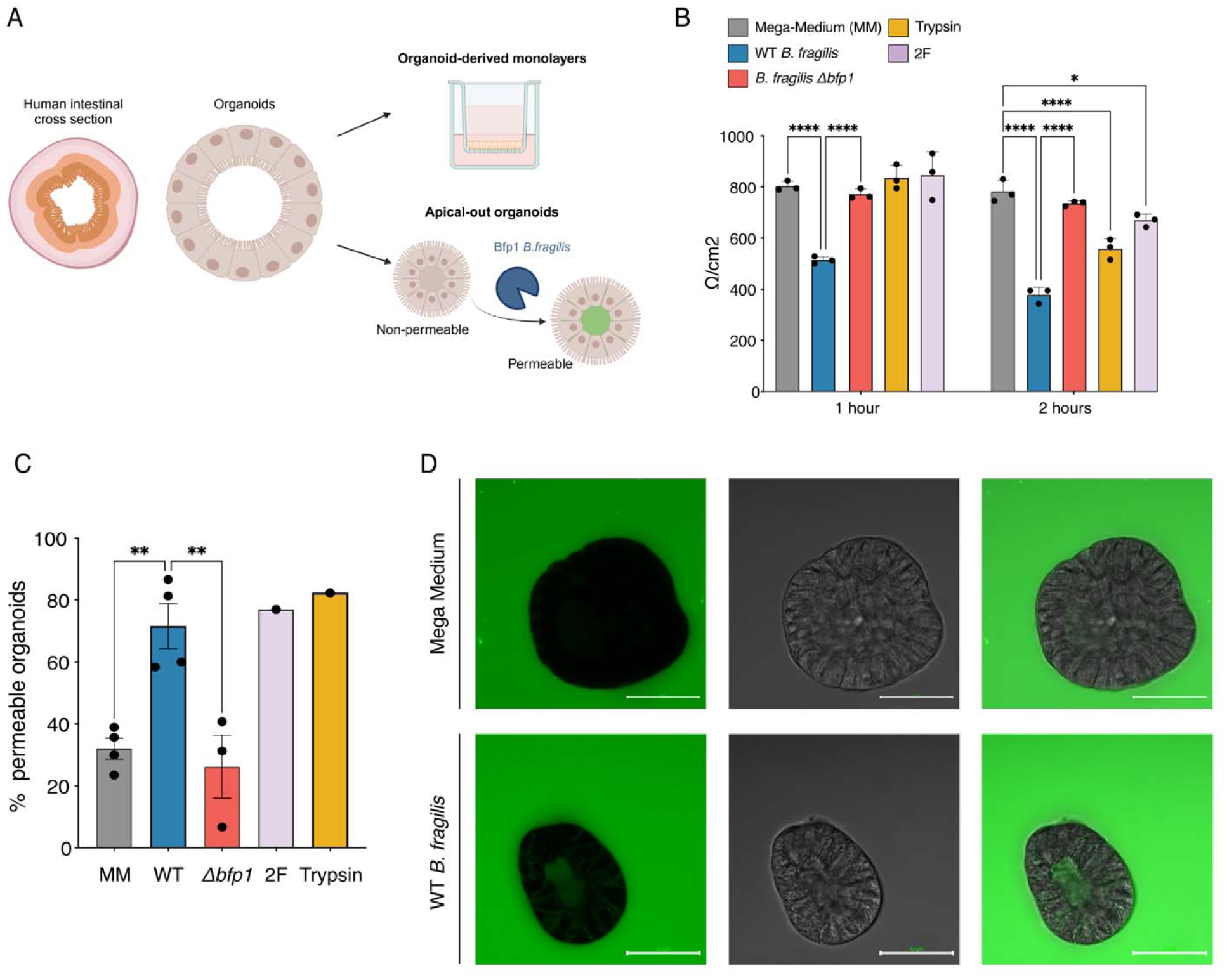
Bfp1 increases epithelial permeability in human intestinal organoids. (A) Experimental workflow for testing epithelial permeability. (B) Time-dependent permeability analysis of organoid-derived monolayers from healthy ileal adult tissue upon exposure to bacterial culture supernatants (4-fold concentrated) or controls, as determined by trans-epithelial electrical resistance (TEER). Trypsin: 4 nM, PAR2-agonist peptide 2-furoyl-LIGRLO-amide (2F): 1 µM, mean + SD, n = 3. (C, D) Permeability analysis of apical-out organoids, as determined by FITC-dextran permeability. Quantitative analysis (C) and representative microscopy images of intact (top) and permeabilized organoids (bottom) (D). Scale bar: 50 µm. Trypsin: 8 nM, 2F: 25 µM, mean + SD, n = 3 or 4 experiments. Each data point represents a percentage determined in a given experiment with 13 – 30 organoids each. Statistical testing via two-way ANOVA. * *p*<0.05, ** *p*<0.01, **** *p*<0.0001. Nonsignificant (ns) test differences are not presented.

Treatment of monolayers with bacterial supernatants resulted in a significantly decreased transepithelial electrical resistance (TEER) for WT *B. fragilis,* but not for *B. fragilis* Δ*bfp1*, which was not significantly different from the MM-treated control (**Figure 4B**). Upon incubation for 2 hours, the residual resistance for WT *B. fragilis*-treated monolayers was 378 ± 30 Ω/cm^2^, which corresponds to 49% of the resistance for *B. fragilis* Δ*bfp1* (736 ± 10 Ω/cm^2^, *p*<0.001). In accordance with these results, culture supernatants from WT *B. fragilis,* but not from *B. fragilis* Δ*bfp1* induced a high percentage of organoids (71.6 ± 7.3 compared to 39.3 ± 10.7, *p*<0.01) that were permeable to FITC-dextran (**Figure 4C**), as determined by confocal microscopy (**Figure 4D**). These findings highlight that Bfp1-containing culture supernatants induce an overall impairment of the epithelial barrier that, during homeostatic conditions, protects the mucosa from inflammatory factors in the intestinal lumen.

### Bfp1 causes PAR_2_-dependent excitation of nociceptors and enhances mechanosensitivity

PAR_2_ is expressed by a subpopulation of dorsal root ganglia (DRG) nociceptors, where activation results in sensitization that leads to pain.^3,31,32^ To determine whether *B. fragilis* proteases also cause PAR_2_-dependent sensitization of nociceptors, we recorded patch clamp measurements of DRG neurons from WT *Par_2_^+/+^* or global *Par_2_^-/-^*knockout (KO) mice. The rheobase (minimum input current required to fire an action potential) of DRG nociceptors was measured to assess sensitization. The rheobase of nociceptors from *Par_2_^+/+^* mice incubated with culture MM (control) for 10 min at RT was 37.6 ± 4.3 pA (n = 29 neurons) (**Figure 5A**). Incubation with WT *B. fragilis* supernatant reduced the rheobase to 15.5 ±1.3 pA (n = 31 neurons, *p*<0.001 to control), denoting heightened excitability. In sharp contrast, supernatant from *B. fragilis* Δ*bfp1* did not affect rheobase. Preincubation of supernatant from WT *B. fragilis* with the serine protease inhibitor FP-alkyne fully abrogated the reduction in rheobase (38.6 ± 2.5 pA, n = 28 neurons, *p*<0.001 to WT supernatant). Similarly, preincubation of neurons with the PAR_2_ antagonist AZ3451^33^ prevented the reduction in rheobase (42.9 ± 5.1 pA, n = 24 neurons, *p*<0.001 to WT supernatant). Corroborating these results, WT *B. fragilis* did not reduce the rheobase of DRG neurons isolated from *Par_2_^-/-^*global KO mice (44.7 ± 9 pA, n = 19 neurons, *p*<0.001 to WT supernatant in WT mice). The results show that WT *B. fragilis* supernatants cause hyperexcitability of mouse nociceptors and that effects are mediated by Bfp1-catalyzed cleavage of PAR_2_.

**Figure 5.**
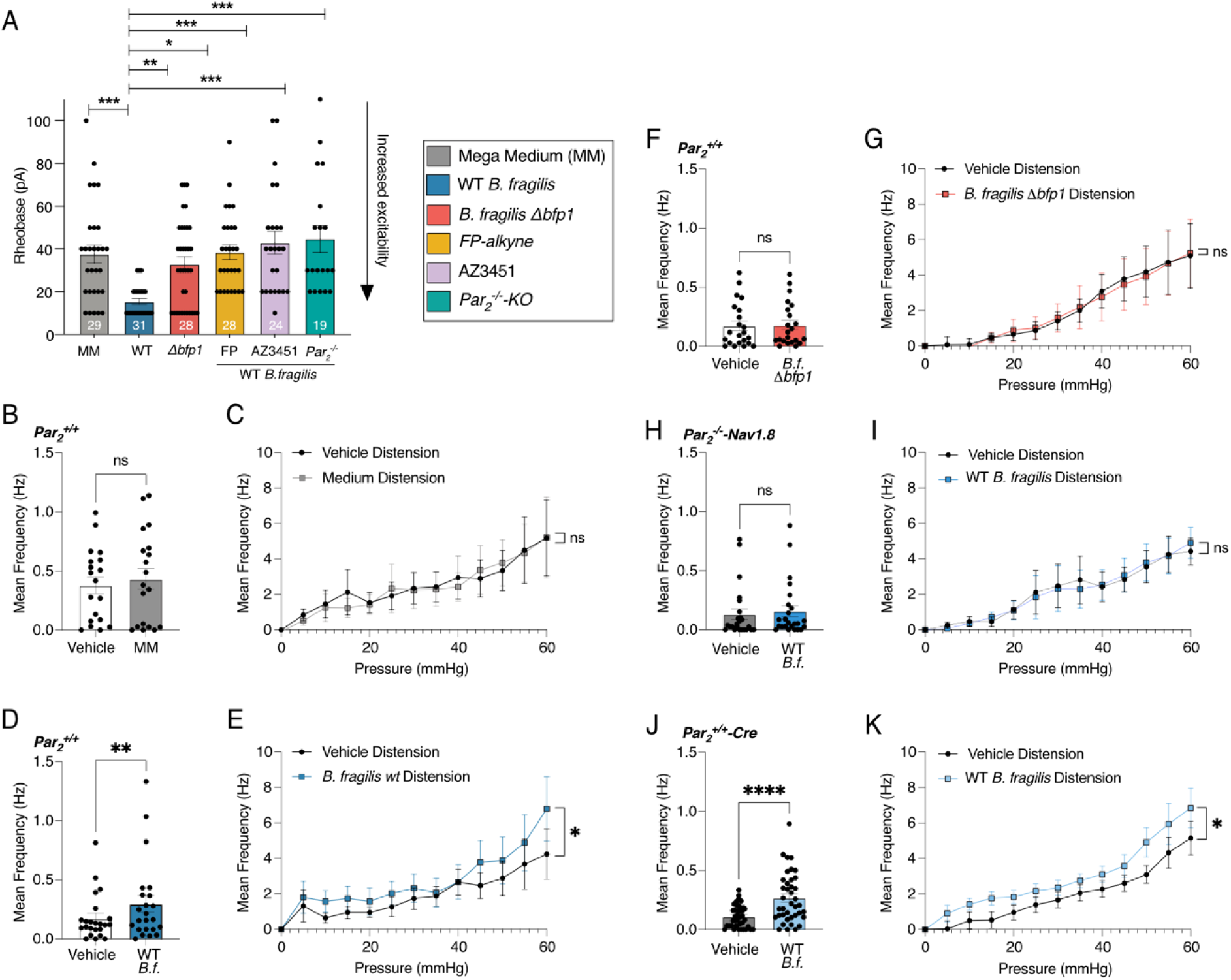
Bfp1 induces nociception and enhances mechanosensitivity in *ex vivo* samples. (A) Excitation of dorsal root ganglia (DRG) nociceptors by 10-fold concentrated *B. fragilis* culture supernatants or Mega Medium (MM) control. To validate Bfp1- and PAR_2_-mediated response, supernatant was pre-treatment with serine protease inhibitor FP-alkyne (100 µM, 60 min, RT), DRG were preincubated with PAR_2_-antagonist AZ3451 (1 µM, 60 min, RT), or neurons from *Par_2_^-/-^* global KO mice were studied. Data show summary of all rheobase activity measurements. Statistical analysis: one-way ANOVA * *p*<0.05, ** *p*<0.01, *** *p*<0.001. Sample numbers are depicted in the plot. (B–K) Effect of supernatants from WT *B. fragilis* and Δ*bfp1* cultures on murine colonic afferent neuron activity in *Par_2_^+/+^*WT, *Par_2_^-/-^*-Nav1.8 KO or *Par_2_^+/+^-Cre* mice. Spontaneous baseline activities (B, D, F, H, J) and overall mechanosensitive response of colonic afferent nerves to distension (C, E, G, I, K). Data are presented as mean[±[SEM. Statistical testing via Wilcoxon test (baseline activity) and two-way ANOVA (mechanosensitive response), ns = not significant, \**p*[<[0.05, \*\**p*[<[0.01, **** *p* < 0.0001.

To determine whether *B. fragilis* proteases can signal from the intestinal lumen to sensitize nociceptor terminals in the colon wall to mechanical stimuli, we performed extracellular recordings for afferent nerves innervating isolated segments of the mouse colon.^32^ Samples were perfused through the colon lumen and action potential firing under baseline conditions and in response to mechanical distension was measured. In WT *Par_2_^+/+^* mice, perfusion of MM alone had no effect on either spontaneous baseline firing (**Figure 5B**, *p* = 0.1594, n = 6 mice, 19 neuronal units) or the response to mechanical distension (**Figure 5C**, *p*□=□0.4217, n□=□6). Culture supernatants from WT *B. fragilis* significantly increased both spontaneous firing (**Figure 5D** and **S6**, *p* = 0.0022, n = 6 mice, 23 neuronal units) and neuronal activation in response to colonic distension (**Figure 5E**, *p* = 0.0406, n = 6). In sharp contrast, perfusion of *B. fragilis* Δ*bfp1* supernatants in *Par_2_^+/+^*colons (**Figure 5F, G**) did not affect spontaneous firing (*p* = 0.4304, n = 6 mice, 22 neuronal units) or the response to colonic distension (*p* = 0.9383, n = 6). Activation of colonic nociceptors and their mechanical sensitization by WT *B. fragilis* supernatants were not observed in colons from *Nav1.8 Par ^-/-^* knockout mice, with targeted deletion of PAR from NaV1.8+ve nociceptors^32^ (**Figure 5H, I**, *p* = 0.0972, n = 6 mice, 24 neuronal units for baseline activity; *p* = 0.9595, n = 6 for distension response). Colons from *Par ^+/+^* Cre-mice (control) showed the same significantly increased responses to culture supernatants from WT *B. fragilis* as observed for WT *Par ^+/+^* mice (**Figure 5J, K**, baseline firing: *p* < 0.0001, n = 7 mice, 39 neuronal units, colonic distension response: *p* = 0.0375, n = 7). Taken together, these results demonstrate that WT *B. fragilis* culture supernatants induce excitation of colonic afferent nerves, which is mediated by Bfp1-triggered activation of PAR_2_ on NaV1.8+ve nociceptors.

### Intracolonic injection of Bfp1 activates PAR_2_ and causes PAR_2_-dependent inflammation and nociception

Mice expressing PAR_2_ C-terminally fused to monomeric ultrastable green fluorescent protein (*Par_2_-mugfp*) allow specific localization of PAR_2_ and analysis of its redistribution during disease.^29^ In the normal colon, PAR_2_ is localized to the basolateral and apical membrane of colonocytes. In the inflamed colon, PAR_2_ redistributes to endosomes of colonocytes, as a result of its proteolytic activation.^29^ It is well established that PAR_2_ signals from endosomes of colonocytes increase paracellular permeability, influx of luminal contents and inflammation, and from endosomes of nociceptors induce hyperexcitability and pain.^29,32^ To assess whether bacterial proteases can activate PAR_2_ in the colon, MM (control) and bacterial supernatants were injected into the colon of *Par_2_-mugfp* mice. After 3 h, the colon was excised and PAR_2_-muGFP was localized by immunofluorescence using GFP antibodies. In mice treated with MM, PAR_2_-muGFP was confined to the basolateral and apical membranes of colonocytes (**Figure 6A**). In mice injected with supernatant from WT *B. fragilis*, PAR_2_-muGFP was depleted from the plasma membrane and detected in endosomes (**Figure 6B**). Supernatant from *B. fragilis* Δ*bfp1* did not cause endocytosis, with PAR_2_-muGFP confined to the plasma membrane (**Figure 6C**). These results are consistent with the hypothesis that Bfp1 from *B. fragilis* cleaves and activates PAR_2_ in the mouse colon, stimulating endocytosis of the receptor.

**Figure 6.**
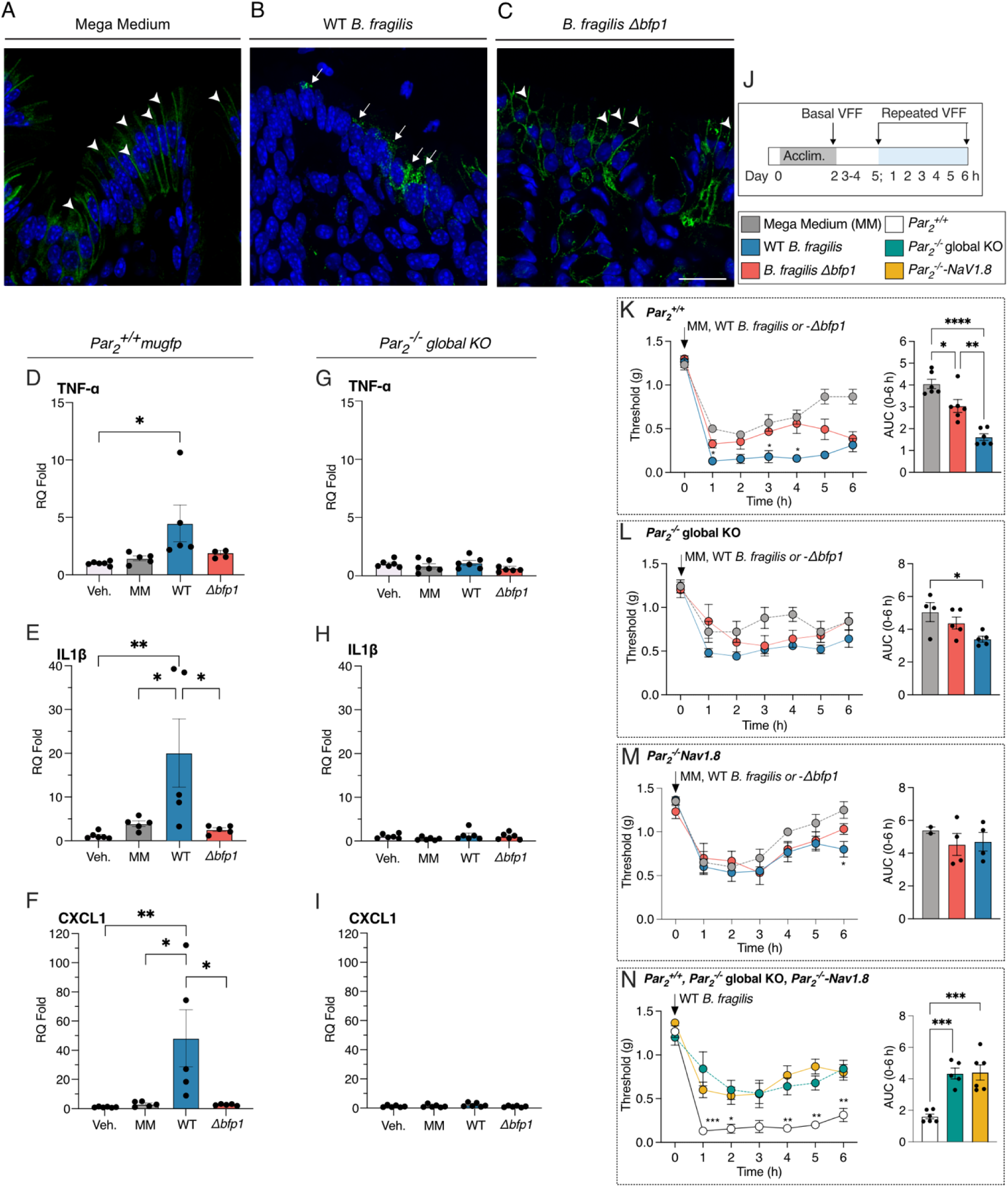
Intracolonic injection of Bfp1 evokes endocytosis of PAR_2_-muGFP, increases proinflammatory cytokines/chemokines and induces colonic nociception through PAR_2_ activation. (A–C) Localization of GFP immunoreactivity in isolated segments of colon from *PAR_2_-muGFP* mice incubated with 10x concentrated (A) Mega Medium (MM), or 10x culture supernatants from (B) WT *B. fragilis* and (C) *B. fragilis* Δ*bfp1*. Arrowheads showing PAR_2_-muGFP localization on the plasma membrane of colonocytes (A, C). Arrows showing PAR_2_-muGFP trafficking in intracellular compartments (B). Representative images, independent experiments, n = 3 mice. Scale bar, 20 µm. (D–I) Colonic cytokine mRNA levels at 3 h after intracolonic administration of Vehicle (Veh.), Mega Medium (MM) and supernatants of WT *B. fragilis* or *B. fragilis* Δ*bfp1* in *Par_2_^+/+^* WT (D–F) and *Par_2_^-/-^* global KO (G–H) mice. Mean ± SEM, One-Way ANOVA, Tukey’s multiple comparison test n = 5 or 6 mice. (J–N) Abdominal mechanical allodynia measurement following intracolonic administration of supernatant (J). Mega medium (MM), *B. fragilis* Δ*bfp1*, and WT *B. fragilis* evoked colonic nociception and area under curve (AUC) in *Par_2_-mugfp* (K)*, Par_2_^-/-^* global *KO* (L) and *Par_2_-Nav1.8* (M). (N) Graphical representation of WT *B. fragilis*-evoked colonic nociception and AUC in *Par_2_-mugfp, Par_2_^-/-^*global *KO,* and *Par_2_-Nav1.8*. Mean ± SEM, n = 5 or 6 mice. Two-way ANOVA, Tukey’s multiple comparison test. \**p*<0.05, \*\**p*<0.01, \*\*\**p*<0.001, \*\*\*\**p*<0.0001.

Notably, intracolonic injection of supernatant from WT *B. fragilis* but not *B. fragilis* Δ*bfp1* stimulated a marked increase in the expression of tumor necrosis factor α (TNFα) (>4-fold higher, *p* = 0.0245), interleukin (IL)-1β (>20-fold higher, *p* = 0.0075) and CXCL1 (>45-fold higher, *p* = 0.0056) mRNA compared to vehicle or MM in the colon of *Par_2_^+/+^-mugfp* mice, consistent with an inflammatory response (**Figure 6D–F**). In contrast, injection of the same samples into the colon of global *Par_2_^-/-^* KO mice did not stimulate any significant cytokine or chemokine transcription (TNFα, 0-fold change, *p* = 0.3184; IL1β, 0-fold change, *p* = 0.3470 and CXCL1, 1-fold higher, *p* = 0.2235; **Figure 6G–I**). Thus, Bfp1 induces PAR_2_-dependent inflammation in the colon.

After intracolonic injection, PAR_2_ agonists are known to cause mechanical allodynia in the colon.^3,29,32^ To determine whether Bfp1 causes mechanical allodynia, we injected MM (control) or bacterial supernatants into the mouse colon. At various time points, we measured withdrawal responses to stimulation of the abdomen with calibrated von Frey filaments (VFF) to assess mechanical allodynia (**Figure 6J**). In WT *Par ^+/+^*mice, Mega Medium (MM) decreased the withdrawal threshold from 1.2 ± 0.0 g (baseline) to 0.5 ± 0.0 g, after 1 h, indicative of mechanical allodynia (**Figure 6K**). WT *B. fragilis* supernatant caused a significantly larger decrease in threshold from 1.3 ± 0.1 g (baseline) to 0.1 ± 0.0 g after 1 h, which was maintained for at least 6 h (0.3 ± 0.1 g), consistent with long-lasting mechanical allodynia of the colon. Supernatant from *B. fragilis* Δ*bfp1* induced a significantly lower response compared to WT supernatant with a baseline threshold of 1.3 ± 0.0 g and a 1 h injection time point mechanical threshold of 0.3 ± 0.1 g (**Figure 6K**). In *Par ^-/-^*global KO mice (**Figure 6L**) or in mice with selective PAR_2_ deletion from Nav1.8+ve nociceptors (*Par ^-/-^Nav1.8*), the withdrawal thresholds to MM, WT *B. fragilis* and *B. fragilis* Δ*bfp1* supernatant were comparable. (**Figure 6M**). Analysis of the area-under-the-curve reveals that the nociceptive threshold changes following the injection of WT *B. fragilis* supernatant in the colon are significantly reduced in *Par* ^+/+^ mice, showing a twofold decrease compared to both *Par ^-/-^* global knockout and *Par ^-/-^Nav1.8* mice (*p* < 0.0001). Thus, the nociceptive response is largely mediated by nociceptor PAR_2_ because it is absent from mice with global or targeted PAR_2_ deletion on Nav1.8+ve nociceptors (**Figure 6N**). Together, these results support the hypothesis that *B. fragilis* Bfp1 activates PAR_2_ in the colon to cause increased sensitivity to mechanical pain.

### Engraftment of the mouse colon with *B. fragilis* triggers abdominal nociception

To determine the impact of secreted Bfp1 on colonic pain in a mouse model, we repopulated antibiotic-treated mice with either WT *B. fragilis* or *B. fragilis* Δ*bfp1* knockout strains (**Figure 7A)**. Mice were treated with an antibiotic cocktail (ampicillin, vancomycin, neomycin, metronidazole) in drinking water for 7 days. We analyzed fecal pellets by qRT-PCR on days 0 (pre-treatment), 3, and 7 of the antibiotic treatment to assess the extent of elimination of colonic bacteria. On day 3, all mice treated with antibiotic cocktail had no detectable bacterial DNA (**Figure S7A**). Mice then received vehicle, WT *B. fragilis* or *B. fragilis* Δ*bfp1* by gavage for 8-19 days to attain engraftment, which was verified by qRT-PCR analysis of fecal pellets for *B. fragilis* DNA (**Figure S7B**). Abdominal nociception was evaluated by measuring withdrawal responses to abdominal stimulation with VFFs. Antibiotic treatment of *Par_2_-mugfp* mice (equivalent to *Par_2_^+/+^* WT)^29^ decreased the threshold for withdrawal to VFF stimulation compared to mice that had not received antibiotics, indicating mechanical allodynia (**Figure 7B-D**). The administration of vehicle to antibiotic-treated mice resulted in a normalization of the withdrawal threshold by 15 days. In marked contrast, in mice repopulated with WT *B. fragilis*, mechanical allodynia was fully sustained during the periods of repopulation and engraftment for at least 19 days (**Figure 7B**). However, repopulation with *B. fragilis* Δ*bfp1* resulted in a partial normalization of mechanical allodynia by day 19. At day 19, the von Frey withdrawal threshold for antibiotic-treated mice was 1.3 ± 0.0 g, 1.2 ± 0.0 g for vehicle-treated mice, 0.3 ± 0.0 g for WT *B. fragilis* repopulated mice, and 0.9 ± 0.1 g for *B. fragilis* Δ*bfp1* repopulated mice (**Figure 7B**). Area-under-the-curve analysis for the period from day 17 to 19 revealed a highly significant (p<0.001) difference between mice engrafted with WT *B. fragilis* (0.3 ± 0.1) compared to mice engrafted with the Δ*bfp1* strain (0.8 ± 0.1, **Figure 7D**). In stark contrast, engraftment of mice with whole-body deletion of PAR_2_ (*Par ^-/-^* global, **Figure 7E–G**) resulted in a drastically reduced colonic nociception upon repopulation with WT *B. fragilis* at day 19 (0.7 ± 0.1 g) that was not significantly different compared to repopulation with *B. fragilis* Δ*bfp1* (0.8 ± 0.1 g, **Figure 7E**). Accordingly, area-under-the-curve analysis for the period from day 8 to 18 (**Figure 7F**) and day 17 to 19 (**Figure 7G**) also did not result in a significant difference between the WT and Δ*bfp1* knockout strains. In contrast to the heightened mechanical allodynia in mice repopulated with WT *B. fragilis*, the mRNA expression of TNFα, IL1β and CXCL1 in the colon at 19 days was not significantly different between antibiotic-treated *Par ^+/+^*or *Par ^-/-^* mice receiving vehicle, WT *B. fragilis* or *B. fragilis* Δ*bfp1* (**Figure S8**).

**Figure 7.**
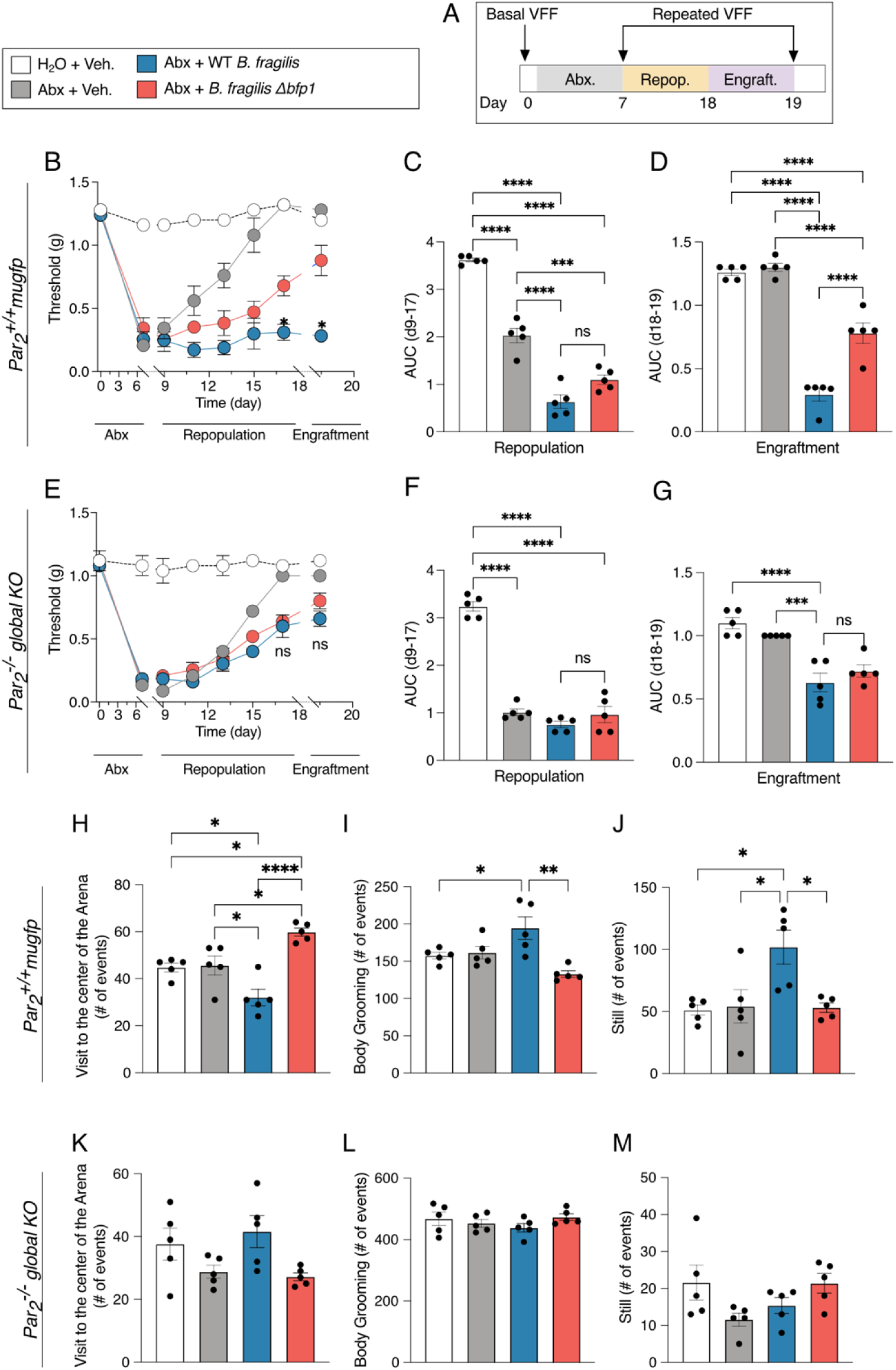
Engraftment of the mouse colon with *B. fragilis* results in Bfp1- and PAR_2_-dependent nociception and behavioral changes. (A–G) Experimental design for the repopulation and engraftment with *B. fragilis* strains after antibiotic depletion of the natural intestinal microbiota of *Par_2_^+/+^* WT (B – D) and *Par_2_^-/-^* -global KO (E–G) mice. (B, E) Abdominal mechanical threshold throughout the whole antibiotics/repopulation experiment. Area under curve (AUC) during repopulation (C, F) and Engraftment (D, G) periods. (H–M) Assessment of spontaneous non-evoked behavior of engrafted mice at day 19 of the experimental protocol. (H, K) Visits to the center of the Arena. (I, L) Body Grooming events, (J, M). Number of still events. Mean ± SEM, n = 5 mice, One-Way ANOVA, and Two-Way ANOVA with Tukey’s multiple comparison test. \**p*<0.05, \*\**p*<0.01, \*\*\**p*<0.001, \*\*\*\**p*<0.0001.

At day 19 of the engraftment period, we evaluated exploratory, grooming and locomotor behaviors of the mice in a behavioral spectrometer, which allows objective quantification of pathological and pain-like behavior in preclinical mouse models of disease (**Figure 7 H–M and S9**).^34^ For *Par_2-_mugfp* mice, there were no detectable differences in any exploratory, grooming or locomotor behavior between non-antibiotic-treated mice and antibiotic-treated mice receiving vehicle or *B. fragilis* Δ*bfp1* (**Figure 7H–J and S9A-C**). In contrast, antibiotic-treated mice receiving WT *B. fragilis* showed significantly decreased activity (**Figure S9A**) and number of visits to the central area compared to mice receiving *B. fragilis* Δ*bfp1* (**Figure 7H**, *p*<0.0001 and **Figure S10, Video 1**). Moreover, antibiotic-treated mice receiving WT *B. fragilis* showed a significant increase in the number of total body grooming events (**Figure 7I***, p*<0.01) and still events (**Figure 7J***, p*<0.05) compared to mice receiving *B. fragilis* Δ*bfp1*. Notably, none of the described exploratory, grooming or locomotor behaviors were significantly different for *Par ^-/-^* global KO mice (**Figure 7K–M and Figure S9E–F**). Taken together, these results are consistent with the hypothesis that *B. fragilis* Bfp1 mediates not only the induction of intestinal permeability and initiation of the pro-inflammatory signaling, but also colonic pain and pain-like behavior in mice *via* the PAR_2_-axis.

## DISCUSSION

The pathogenesis of IBD is multifactorial, with the underlying molecular mechanisms for onset and progression thought to be mainly controlled by three parameters: microbiota, epithelial barrier and immune cells.^35–37^ Previous studies have linked excessive protease activities and elevated PAR_2_- signaling to disrupted barrier function, inflammation, abdominal pain and IBD severity,^11,19,38^ but the role of specific gut bacterial proteases has remained primarily a source of speculation.^9–12,19,39^ Abdominal pain is also a unifying symptom of IBS, and has been related to increased proteolysis and activation of PAR_2_ in the colon.^3,32,40^ To identify bacterial proteases capable of cleaving PAR_2_, we developed an assay that enables direct screening of commensal bacteria found in the human gut for their ability to produce and secrete proteases that cleave PAR_2_. By using the full-length N-terminal domain of PAR_2_, it was possible to screen for proteases that cleave the receptor at any location within the N-terminus accessible to an extracellular enzyme. Using this approach, we found that a substantial number of intestinal bacteria (19% of our 140-member library), particularly *Bacteroides* spp., secrete PAR_2_-processing proteases (**Figure 1C, D**). Subsequent functional studies using broad-spectrum inhibitors and activity-based probes showed that a previously uncharacterized serine protease Bfp1, produced by *Bacteroides fragilis*, cleaves the N-terminus of PAR_2_, resulting in impaired barrier function, pain and intestinal inflammation. These findings identify a potentially valuable new target for therapeutics that could suppress some of the key pathological symptoms of IBD and IBS,^3,41,42^ and highlight the likely complex set of interactions taking place between gut bacteria and the host mediated by secreted microbial proteases. Furthermore, because diverse bacterial species produce protease activities that are capable of cleaving PAR_2_, this receptor could serve as a possible nexus for competition between bacteria in a community.

Cleavage of PAR_2_ by Bfp1 results in receptor endocytosis (**Figure 3**), which is consistent with its canonical activation.^29,32^ However, PARs are GPCRs capable of triggering diverse downstream processes through ‘biased signaling’, wherein the location of proteolytic cleavage determines coupling to specific G-proteins and distinct signaling pathways.^20,43^ Since our assay detects cleavage at any location along the full N-terminal domain, our top hits from the screen have the potential to mediate a diverse range of downstream effects, either through (i) activation by canonical or biased signaling or (ii) receptor inactivation *via* removal of the tethered ligand. The ability to modulate multiple activities of PAR_2_ may markedly affect gut physiology, considering the central role of PAR_2_ in controlling epithelial barrier integrity, colonic inflammation and visceral pain.^32,44–46^ Given that we identified ten putative serine proteases/peptidases in the supernatant of *B. fragilis*, it was striking that genetic deletion of Bfp1 alone was sufficient to disrupt PAR_2_-activation. This suggests that the cohort of proteases secreted by *B. fragilis* and other Bacteroidetes spp. may play different roles in modulating host or microbial processes and that we have only identified one such node which involves PAR_2_ as a substrate.

Previous studies found that antibiotic treatment reduces bacterial serine protease activity in the mouse colon, leading to diminished PAR_2_ processing and expression by colonocytes, but the source of this activity has remained unclear.^47^ Beyond the previously identified PAR_2_ activator from *E. faecalis,*^16^ our work expands the limited list of PAR_2_-NTD-processing bacteria in the human intestine^18,48^ to include *Clostridium sporogenes* and 14 diverse Bacteroidetes species (representing 24 strains). Notably, Bfp1 is one of the few bacterial proteases identified and validated as a PAR_2_-activating enzyme. Although the strong proteolytic activity of *B. fragilis* and other Bacteroidetes spp. was first characterized more than 35 years ago,^49–51^ these bacteria have historically been studied primarily for their ability to degrade complex carbohydrates and produce bioactive metabolites. Despite these advances, the functional roles of Bacteroidetes proteases remain underexplored,^52,53^ with few proteases mechanistically characterized.^54–59^ Notably, the human gut metagenome contains over 285 putative bacterial serine proteases,^8^ making targeted enzyme identification a significant challenge. Using our competitive ABPP approach (**Figure 2**), we efficiently pinpointed Bfp1 as the protease responsible for PAR_2_ proteolysis, demonstrating the power of chemical proteomics to functionally annotate uncharacterized microbial enzymes.

Our finding that Bacteroidetes spp. secrete PAR_2_-processing proteases aligns well with recent reports that demonstrate a link between bacteria-associated proteolysis and IBD/IBS.^60–62^ A recent multi-omics study revealed uncharacterized proteases secreted from *B. vulgatus* that induce colitis in mouse models.^63^ Notably, as *B. vulgatus* is also a top hit in our screen, it is intriguing to hypothesize that the reported proinflammatory phenotypes are mediated by proteolytic processing of PAR_2_.

While enterotoxigenic *B. fragilis* (ETBF), producing the metalloprotease fragilysin (BFT), contributes to acute diarrheal secretory diarrhea and colonic epithelial damage, non-toxigenic *B. fragilis* is generally regarded as a commensal in the human intestine.^64^ However, studies focused on the role of *B. fragilis* in IBD and IBS have yielded diverse outcomes, with evidence for both protective^65–67^ and pathogenic^68–71^ effects. The majority of these studies focus on metagenomic analyses which are often correlative. In the current study, we identify a specific bacterial effector, Bfp1, its human receptor, PAR_2_, and the physiological phenotypes (loss of barrier function, inflammation and pain) that result from their interaction, thus providing a mechanistic understanding of this specific axis of host-microbe interactions. The seemingly conflicting results of prior studies and our findings may, in part, reflect differences in the regulation and expression of secreted factors like Bfp1, which could be influenced by presently unknown molecular cues or environmental stimuli. Furthermore, *B. fragilis* may have evolved Bfp1 to activate PAR_2_-signaling and create an ecological niche conducive to its survival. Conversely, other PAR_2_-processing activities identified in our proteolysis screen might exert opposing effects to those of Bfp1, potentially deactivating PAR_2_-signaling as a strategy to suppress inflammatory conditions or to counteract dominance of a species such as *B. fragilis*. PAR_2_ activation can induce both pathologic and protective effects, which may also account for the detrimental and beneficial actions of *B. fragilis*.^20^

While there are a number of animal models of IBD, current advances in the use of intestinal organoids enable establishment of highly controllable cell models that mimic the interactions between microbes and human gut tissue. Previous work using classical organoids in which *B. fragilis* supernatants were added basolaterally suggested that *B. fragilis* does not impact barrier integrity.^70^ In contrast, we find that physiologically-relevant apical treatment of organoids with supernatant containing Bfp1 disrupts barrier function to a similar extent as an optimized PAR_2_ agonist or trypsin, a protease known to activate PAR_2_ (**Figure 4**). This result demonstrates the value of the apical-out organoid model^30^ and suggests that the prior inability to detect an impact of *B. fragilis* on barrier function may be attributed to the inability of secreted proteases to reach the epithelial apical side. A loss of barrier function triggered by Bfp1 and PAR_2_ would be expected to induce an influx of bacteria and macromolecules from the colon lumen, leading to colonic inflammation and pain.

Our findings that Bfp1 increases excitability of isolated DRG nociceptors, sensitizes colonic nociceptors to mechanical stimuli, and evokes abdominal mechanical allodynia and pain-like behavior in mice through mechanisms largely dependent on PAR_2_ are in line with the known actions of PAR_2_ in the colon. PAR_2_ agonists, including proteases and peptide analogs of the tethered ligand, sensitize transient receptor potential ion channels^72^ and induce hyperexcitability^32^ of nociceptors. Moreover, proteases from macrophages, mast cells, and from colon biopsies of IBS patients induce visceral nociception upon intracolonic injection by a PAR_2_-mediated mechanism.^3,29,32,46,73^ PAR_2_ agonists also cause a loss of barrier function in the colon^29,45^ and stimulate the release of neuropeptides from colonic nociceptors,^44,74^ which would lead to inflammation and ensuing pain. Activation of PAR_2_ in nociceptors and colonocytes would be expected to cause pain in mice. However, our results that selective deletion of PAR_2_ from NaV1.8 nociceptors diminishes the pronociceptive actions of Bfp1 to a similar extent as global PAR_2_ deletion, suggests that Bfp1 mostly evokes pain through nociceptor-PAR_2_.

Collectively, our findings that Bfp1-positive culture supernatants consistently induce PAR_2_-dependent phenotypes across organoid, neuronal, and mouse models, closely reflecting the established effects of PAR_2_ signaling, provide strong evidence for a physiologically relevant axis of host-microbe interaction. By identifying Bfp1 as a molecular effector of intestinal pain, barrier function and inflammation, this study establishes a crucial framework to investigate how microbial proteases drive pathogenesis. Future research will be required to build upon these findings to further elucidate the complex roles of microbial proteases and explore their potential as therapeutic targets for restoring intestinal homeostasis and mitigating inflammatory and painful diseases.

### Limitations of the study

1. While our data shows that Bfp1 processes and activates PAR_2_ to mediate nociceptive and inflammatory phenotypes, there may be additional substrates for Bfp1 that are important for the regulation of other biological processes (e.g. activation of other PARs or degradation of tight-junction proteins). Future studies will be required to map the substrates of Bfp1 in both bacterial and human proteomes.
2. While our data shows that Bfp1 alone is able to trigger PAR_2_ signaling, it is not clear if it acts as a dominant regulator of the pathogenesis of IBD or IBS. More detailed studies will be required to determine if inhibition of Bfp1 is sufficient to produce a protective effect in IBD/IBS models and then eventually whether Bfp1 inhibitors can be used as a strategy to treat human patients with these diseases.
3. Our study has identified multiple bacterial strains from the human gut microbiome that secrete enzymes that can cleave the PAR_2_ N-terminal domain at some location. While we identified Bfp1 as a novel protease that can cleave PAR_2_, future work will be needed to characterize the additional PAR_2_ processing proteases to begin to map the complex network of proteolytic interactions within the intestine.

## ACKNOWLEDGEMENTS

This work was supported by NIH grant R01 DK130293 (to M.B. and NB), R01 NS125413 (to D.J.) and funding from Takeda Pharmaceuticals (to M.B. and N.B.). M.L. thanks the Deutsche Forschungsgemeinschaft (DFG, German Research Foundation) for funding via the Walter-Benjamin-Fellowship (project ID 450273105) and the *Fonds der Chemischen Industrie* for funding via their Liebig-Fellowship. N.S. and M.L are grateful for financial support by the research profile LIFE of Friedrich-Schiller University Jena and the Deutsche Forschungsgemeinschaft (DFG, German Research Foundation) via the Emmy-Noether-Program (project ID 528114058) and Germanýs Excellence Strategy (EXC 2051, project ID 390713860). A.L., D.R. and H.W. gratefully acknowledge funding from the Canadian Institutes of Health Research (project 166053). K.B. was supported by a Postdoctoral fellowship from the Stanford Medicine Children’s Health Center for IBD and Celiac Disease. L.J.K. was supported by the Stanford ChEM-H Chemistry/Biology Interface Predoctoral Training Program (T32 GM120007), a Stanford Molecular Pharmacology Training Grant (T32 GM113854), and a Stanford Graduate Fellowship. We are grateful to Will van Treuren, Shuo Han and Justin L. Sonnenburg for providing the bacterial strain library and consulting on bacterial cultivation. We thank Elias Roth Gerrick and Michael R. Howitt for experimental guidance and anaerobic chamber access. We thank Qinghui Mu and Calvin Kuo for providing human ileal organoids and Manuel Amieva for consulting on apical-out organoid culture. This work utilized the Thermo Orbitrap Eclipse nanoLC/MS system (RRID:SCR_022212) that was purchased with funding from National Institutes of Health Shared Instrumentation Grant 1S10OD030473.

## Supplemental Figures

**Figure S1.**
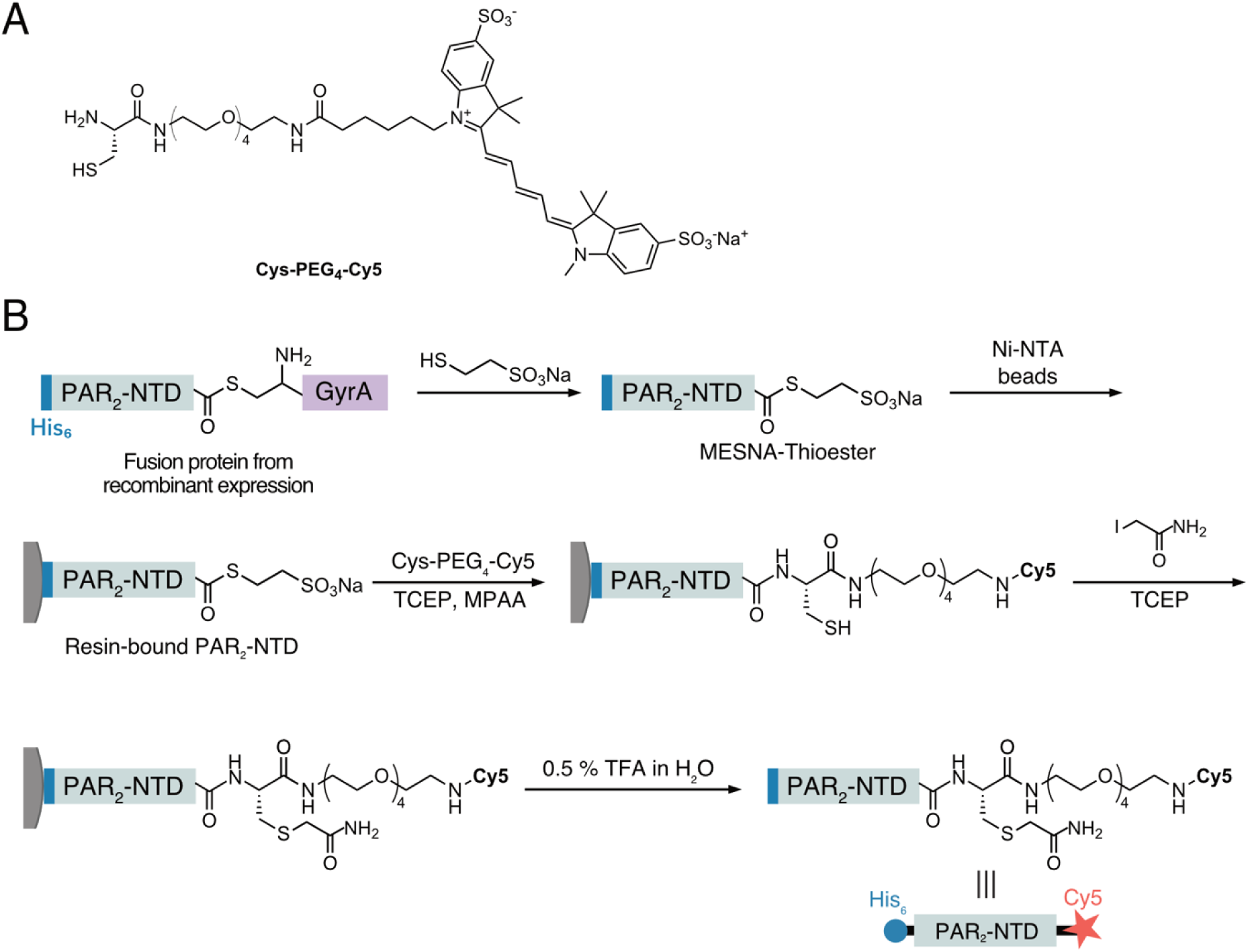
Generation of dual-labeled substrate His_6_-PAR_2_NTD-Cy5 by expressed protein ligation. (A) Chemical structure Cys-PEG_4_-Cy5. See Supplementary Note 1 for its chemical synthesis. (B) Schematic workflow for on-resin native chemical ligation of His_6_-PAR_2_-NTD-GyrA and Cys-PEG_4_-Cy5 to generate the dual-labeled substrate.

**Figure S2.**
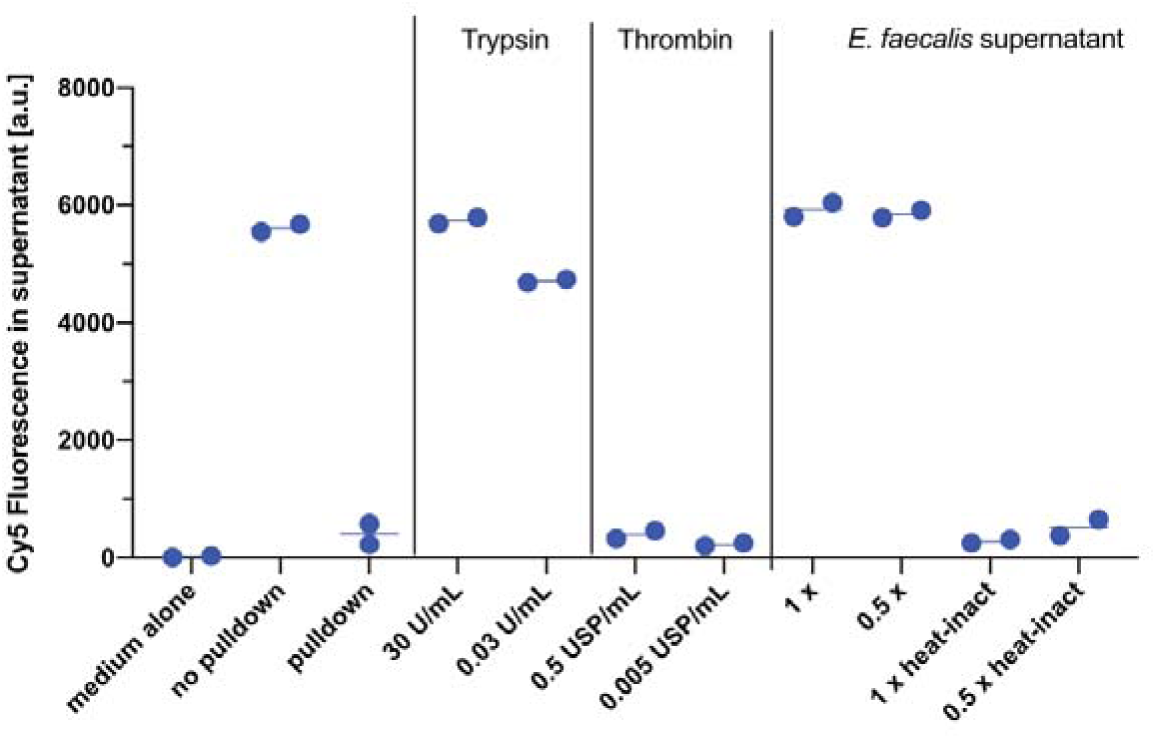
Establishment and optimization of the PAR2-proteolysis assay. Trypsin and culture supernatant from *E. faecalis* cultures, containing the protease gelatinase, were positive controls. Thrombin and heat-inactivated *E. faecalis* culture supernatant served as negative controls.

**Figure S3.**
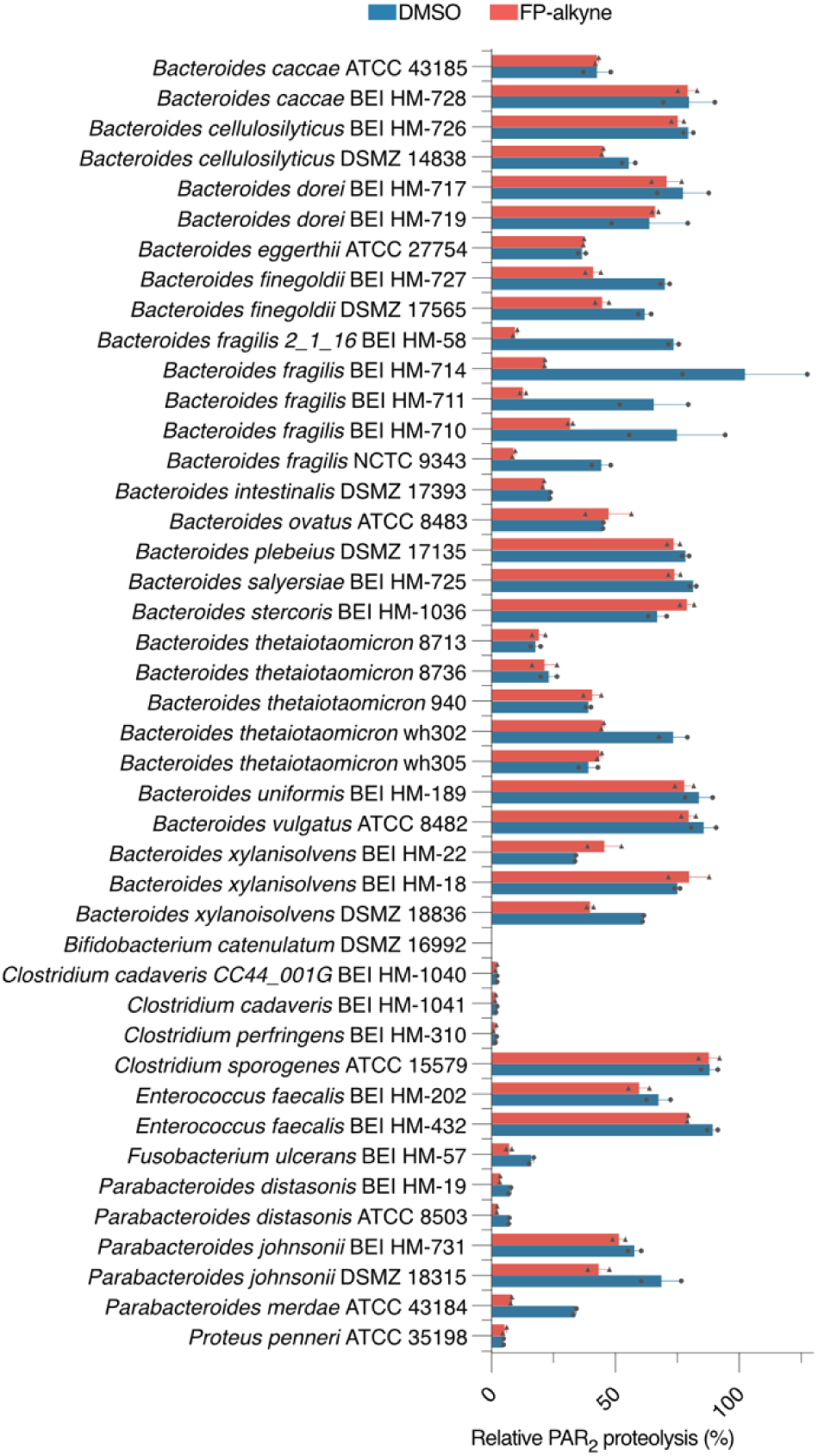
Validation of PAR2-proteolytic activity. Relative activities of all original “hit” supernatants harvested at early stationary phase. Activities upon pre-incubation with DMSO (control) or FP-alkyne (20 µM) for 1 h. Proteolysis duration: 1.5 h (instead of 3 h for the initial screen). Data are normalized to treatment with trypsin (100% processing) and presented as mean ± SEM, n = 2.

**Figure S4.**
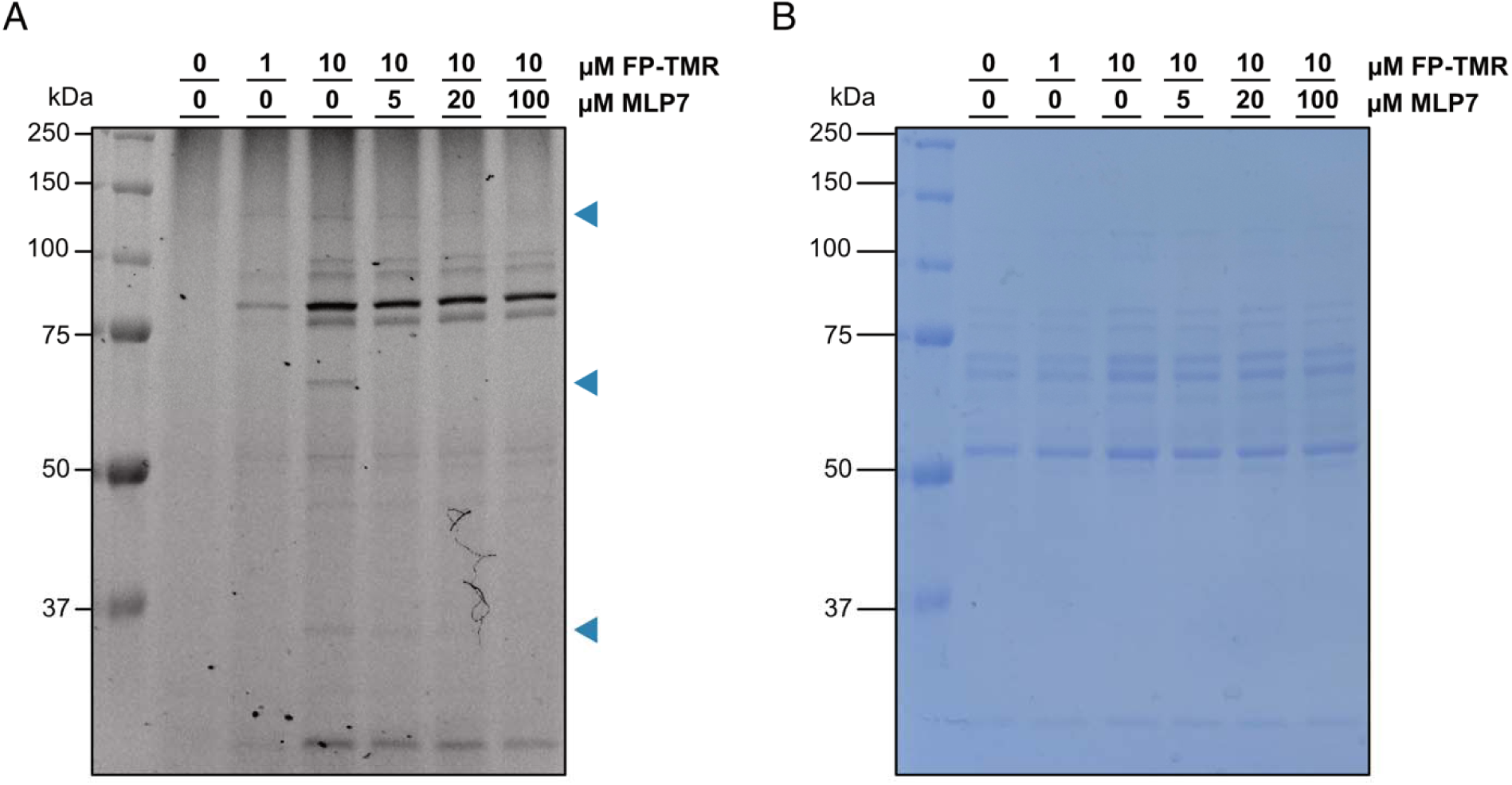
Gel-based ABPP for optimization of labeling conditions. (A) Fluorescent scan of SDS-PAGE gel of *B. fragilis* NCTC 9343 culture supernatants competitively labeled with **MLP7** and visualized by **FP-TMR**. (B) Corresponding Coomassie-stained SDS-PAGE gel.

**Figure S5.**
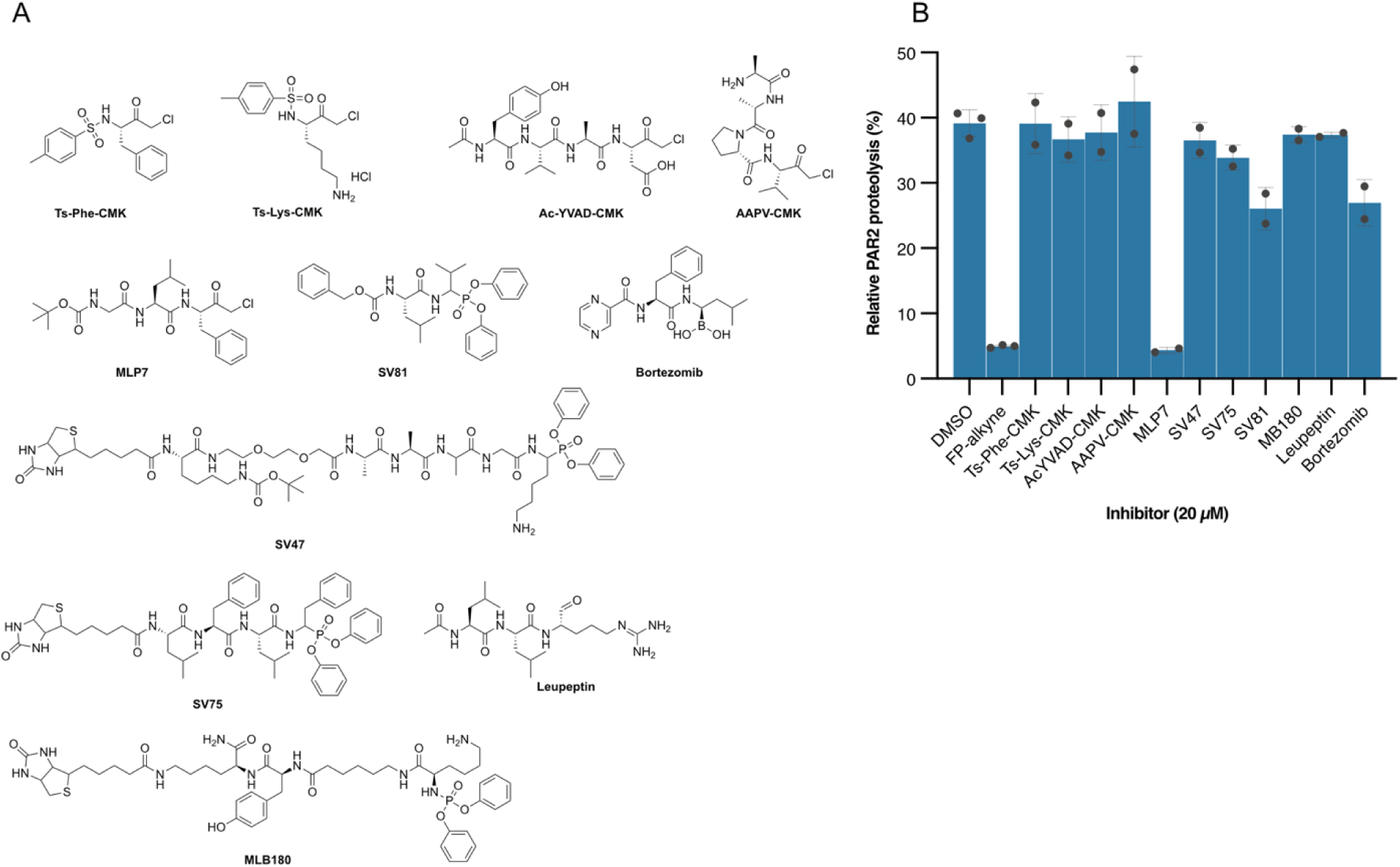
Screening of covalent inhibitors that inhibit PAR_2_-processing of the *B. fragilis*. (A) Chemical structures of inhibitors tested. (B) Relative inhibition (normalized to 100% processing) of the proteolysis of the PAR_2_-NTD substrate by of *B. fragilis* culture supernatants. Mean ± SEM, n = 3 or 2.

**Figure S6:**
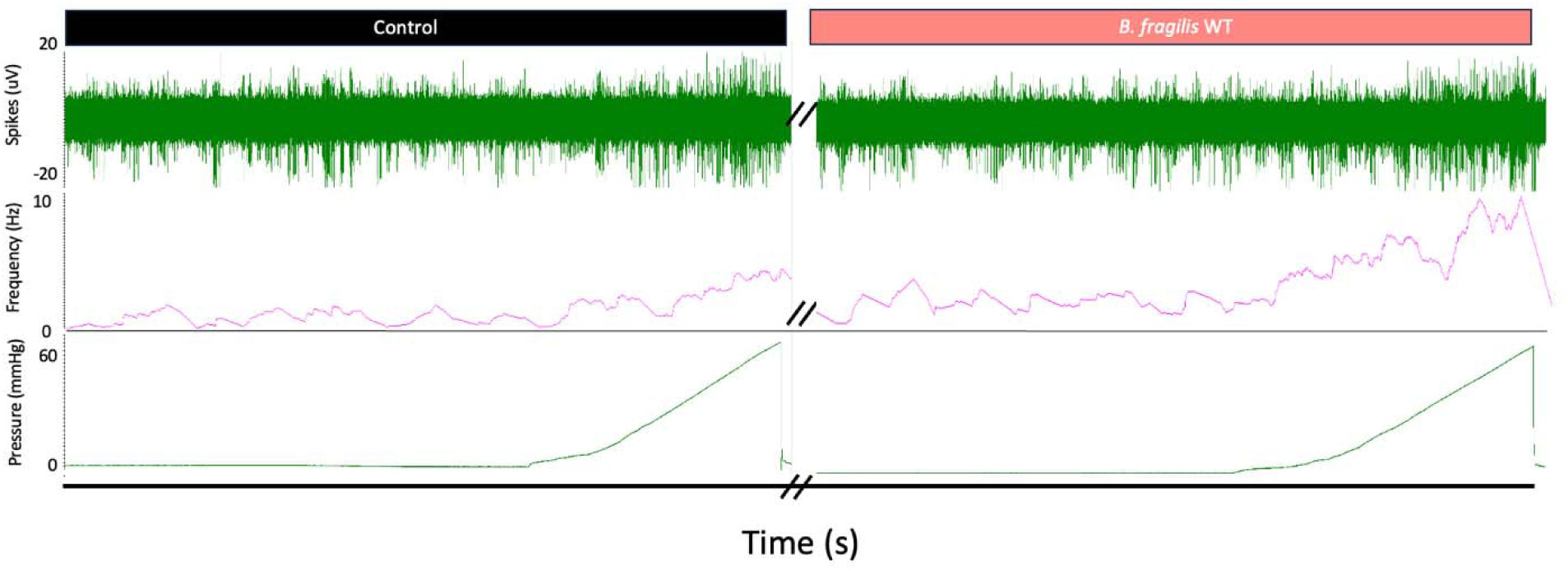
Representative afferent nerve recording trace of WT *B. fragilis* perfusion. *Left* side depicts the control baseline activity and ramp distension during Krebs vehicle perfusion, while the *right* depicts the increase in baseline activity frequency and mechanical response to ramp distension during perfusion of WT *B. fragilis*.

**Figure S7.**
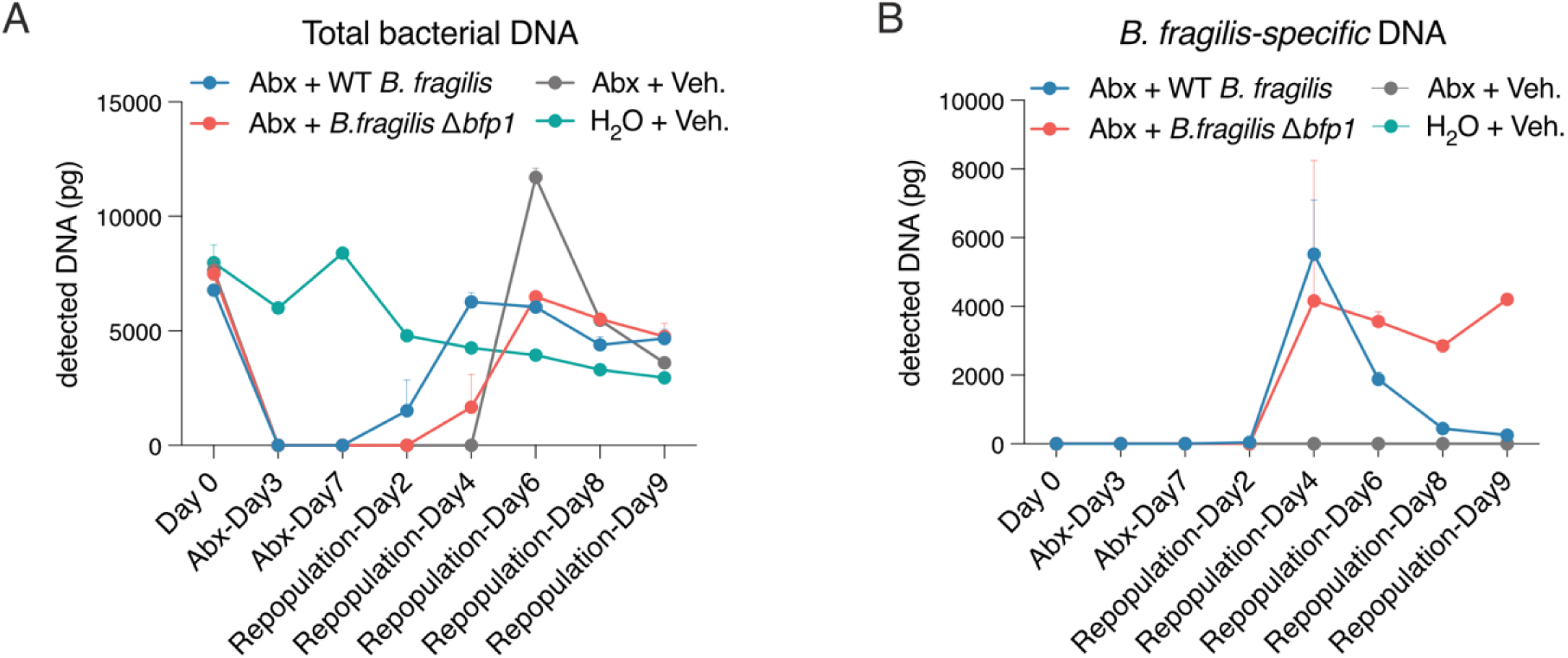
Validation of bacterial deletion and repopulation with *B. fragilis*. (A) qRT-PCR analysis for total bacterial DNA found in the feces of mice. (B) qRT-PCR analysis for *B. fragilis*-specific DNA found in the feces of mice. A, B: Mean ± SEM, n = 3 or 4.

**Figure S8.**
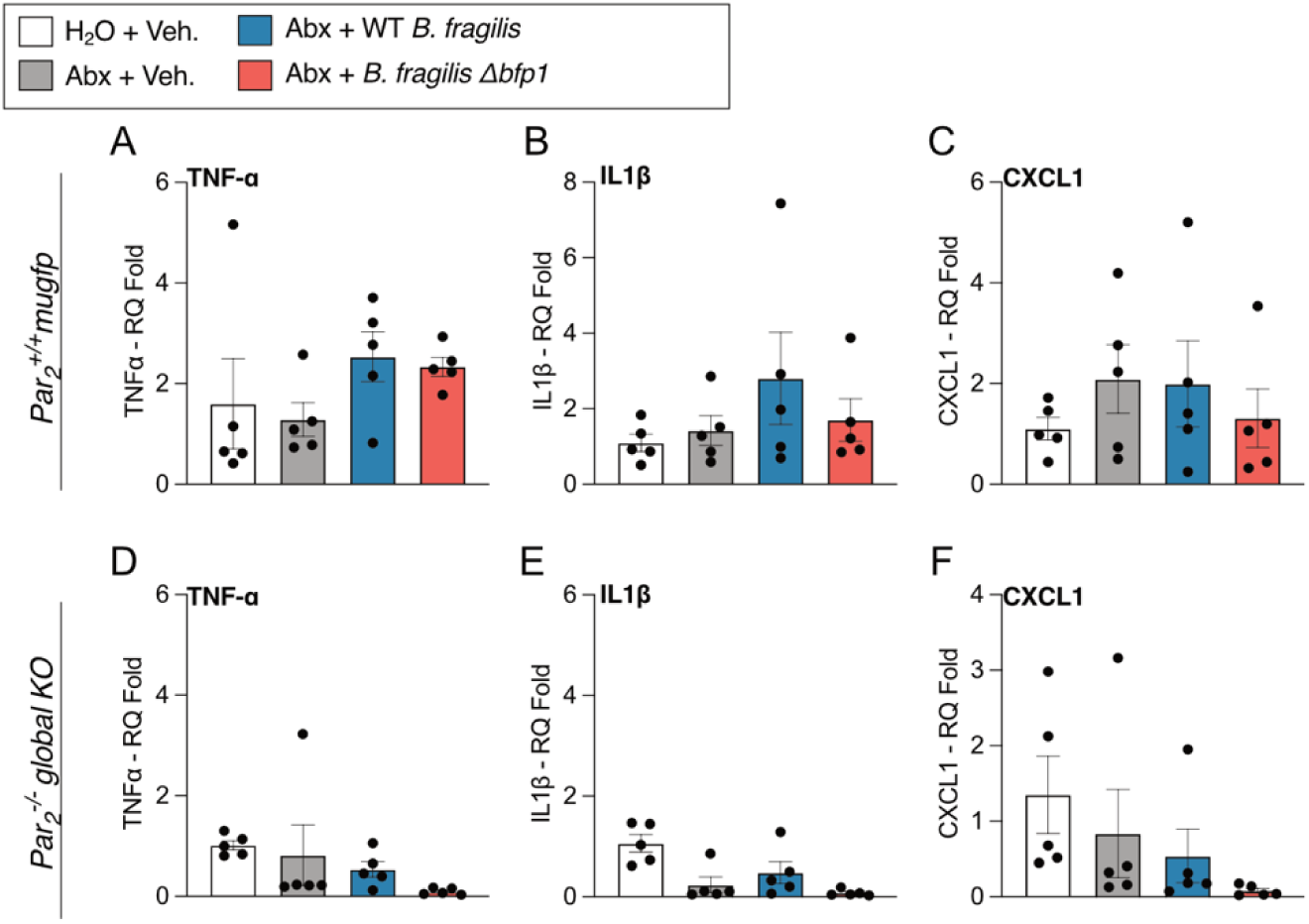
Colonic cytokine/chemokine mRNA levels in mice following bacterial repopulation. (A–F) Colonic cytokine/chemokine mRNA levels of *Par_2_^+/+^* mugfp mice (A–C) and *Par_2_^-/-^* global knockout mice (D–F) at day 19 of the repopulation experiment. Mean ± SEM, One-Way ANOVA, Tukey’s multiple comparison test n = 5 mice. \**p*<0.05, \*\**p*<0.01, \*\*\**p*<0.001, \*\*\*\**p*<0.0001.

**Figure S9.**
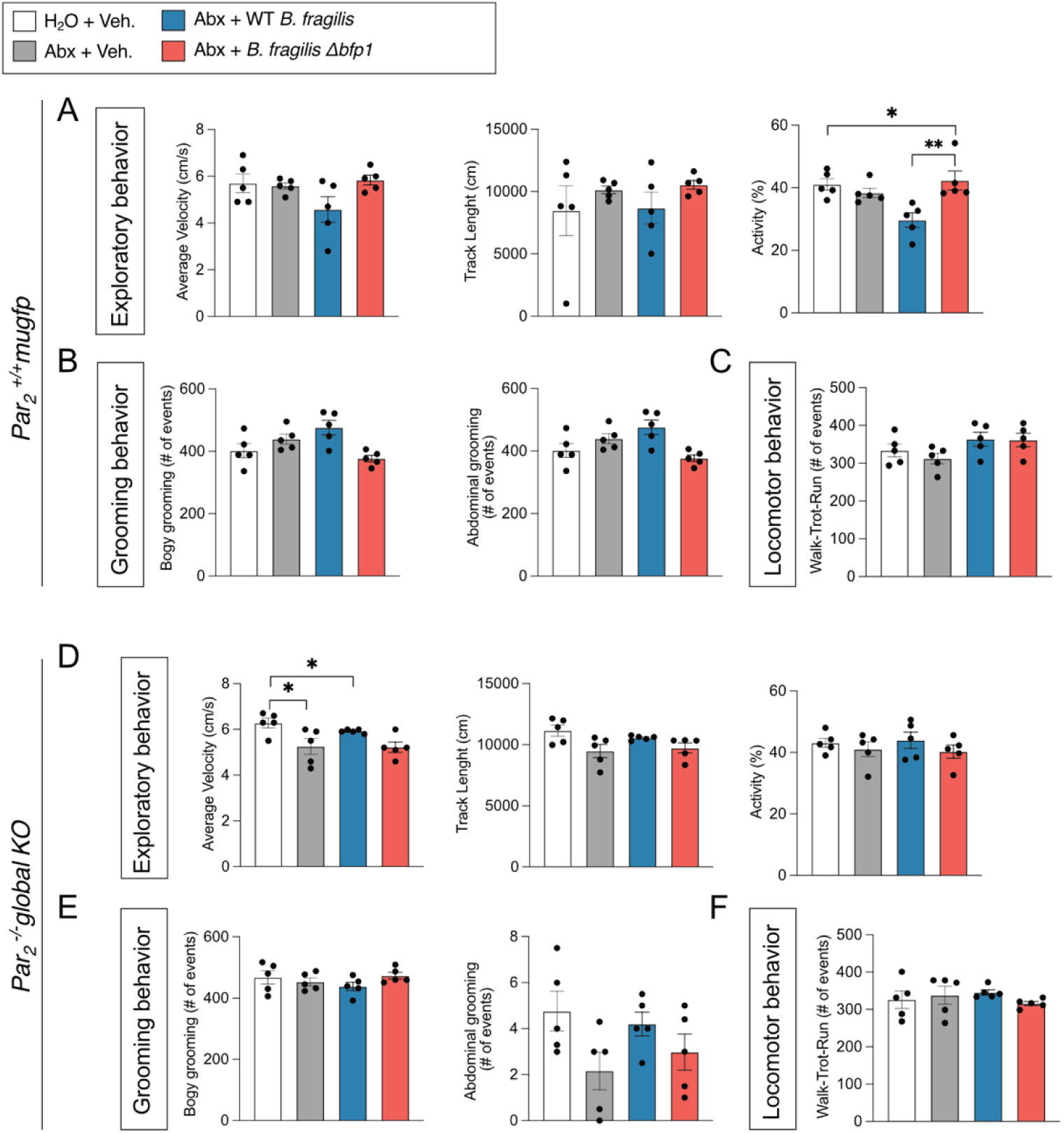
Spontaneous behavior assessment in mice following bacterial repopulation. (A–C) Behavior of *Par_2_^+/+^* mugfp mice on day 19 (engraftment). (A) Exploratory behavior (average velocity, track length, activity), (B) Grooming behavior (number of body grooming events, number of abdominal grooming events), (C) Locomotive behavior (number of walk-to-run events). (D–F) Behavior of *Par_2_^-/-^*global knockout mice on day 19 (engraftment). (D) Exploratory behavior (average velocity, track length, activity), (E) Grooming behavior (number of body grooming events, number of abdominal grooming events), (F) Locomotive behavior (number of walk-to-run events). For (A-F): Behavior was monitored for 30 min using a behavioral spectrometer, mean ± SEM, n = 5 mice. * *p*<0.05, ** *p*<0.01. One-way ANOVA, Tukey’s multiple comparison test.

**Figure S10.**
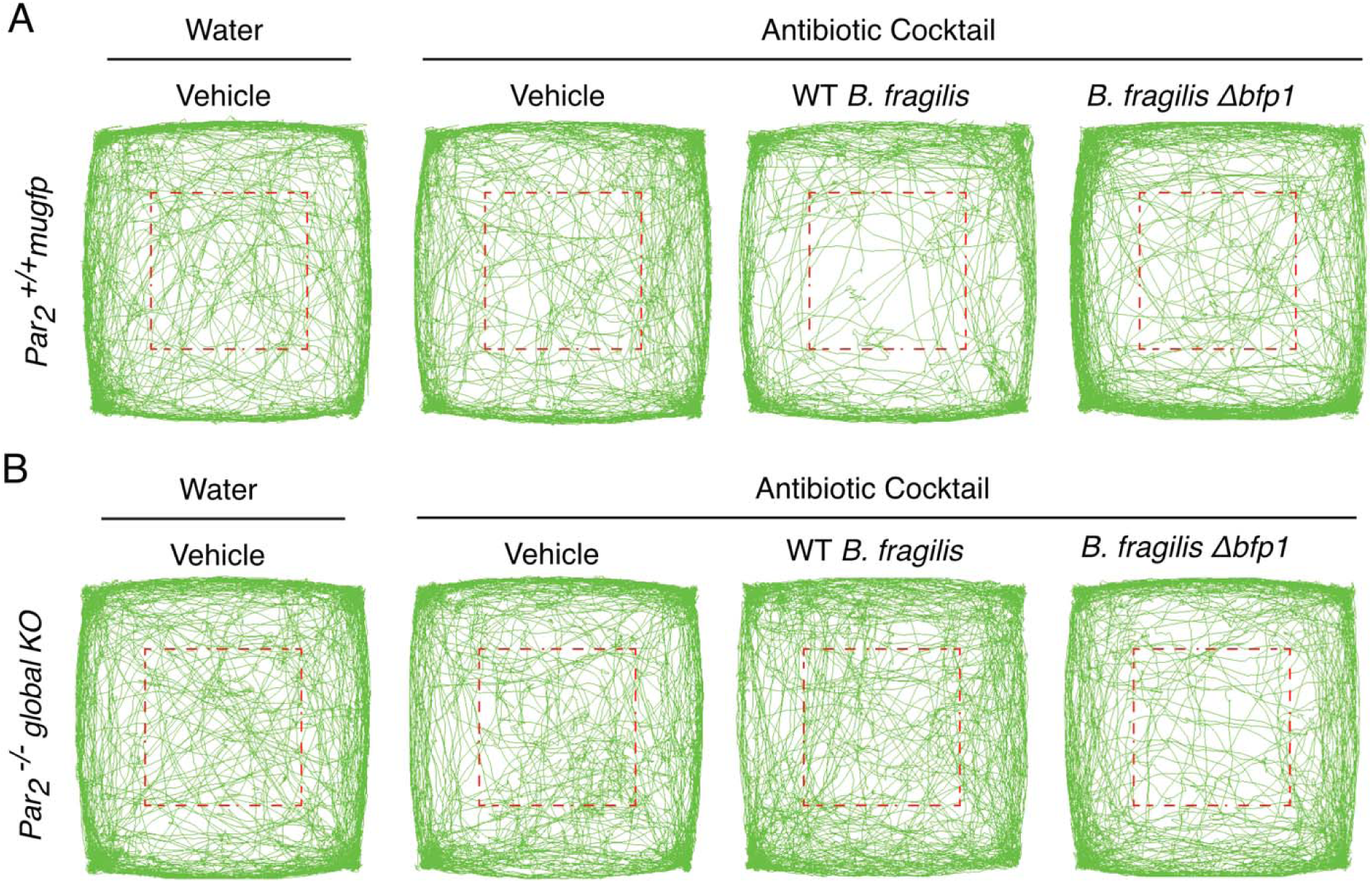
Spontaneous behavior of mice after antibiotic treatment, repopulation and engraftment with *B. fragilis*. (A–B) Traces show the spontaneous behavior of mice measured over 30 min in a behavioral spectrometer on day 19 of the antibiotic treatment, repopulation and engraftment protocol. (A) *Par ^+/+^ mugfp* mice and (B) global *Par ^-/-^* mice received water and vehicle (control), or antibiotic cocktail followed by vehicle, WT *B. fragilis* bacteria, or *B. fragilis* Δ*bfp1* bacteria. The green lines record the movement of n = 5 mice. The red dashed line shows the central area.

**Video 1. Spontaneous behavior of mice after antibiotic treatment, repopulation and engraftment with *B. fragilis*.** The videos show the spontaneous behavior of mice over 90 s (speeded-up three times) on day 19 of the antibiotic treatment, repopulation and engraftment protocol. *Par_2_^+/+^ mugfp* mice received water and vehicle (control) or antibiotic cocktail followed by vehicle, WT *B. fragilis*, or *B. fragilis* Δ*bfp1*. Recordings are from individual mice that are representative of n = 5 mice.

## REFERENCES

1. Azlan, P.M.M., Chin, S., Low, T.Y., Neoh, H., and Jamal, R. (2019). Analyzing the Secretome of Gut Microbiota as the Next Strategy For Early Detection of Colorectal Cancer. Proteomics, 1800176. 10.1002/pmic.201800176.

2. Hildner, K., Waldschmitt, N., and Haller, D. (2018). Microbiome and Diseases: Inflammatory Bowel Diseases. In The Gut Microbiome in Health and Disease, D. Haller, ed. (Springer, Cham.), pp. 151–174. 10.1007/978-3-319-90545-7_11.

3. Cenac, N., Andrews, C.N., Holzhausen, M., Chapman, K., Cottrell, G., Andrade-Gordon, P., Steinhoff, M., Barbara, G., Beck, P., Bunnett, N.W., et al. (2007). Role for protease activity in visceral pain in irritable bowel syndrome. J. Clin. Investig. 117, 636–647. 10.1172/jci29255.

4. Cryan, J.F., O’Riordan, K.J., Sandhu, K., Peterson, V., and Dinan, T.G. (2020). The gut microbiome in neurological disorders. Lancet Neurology 19, 179–194. 10.1016/s1474-4422(19)30356-4.

5. Huang, J., Liu, W., Kang, W., He, Y., Yang, R., Mou, X., and Zhao, W. (2022). Effects of microbiota on anticancer drugs: Current knowledge and potential applications. EBioMedicine 83, 104197. 10.1016/j.ebiom.2022.104197.

6. Lee, W.-J., and Hase, K. (2014). Gut microbiota-generated metabolites in animal health and disease. Nat. Chem. Biol. 10, 416–424. 10.1038/nchembio.1535.

7. Donia, M.S., and Fischbach, M.A. (2015). Small molecules from the human microbiota. Science 349, 1254766. 10.1126/science.1254766.

8. Kriaa, A., Jablaoui, A., Mkaouar, H., Akermi, N., Maguin, E., and Rhimi, M. (2020). Serine proteases at the cutting edge of IBD: Focus on gastrointestinal inflammation. Faseb J. 10.1096/fj.202000031rr.

9. Vergnolle, N. (2016). Protease inhibition as new therapeutic strategy for GI diseases. Gut 65, 1215– 1224. 10.1136/gutjnl-2015-309147.

10. Spaendonk, H.V., Ceuleers, H., Witters, L., Patteet, E., Joossens, J., Augustyns, K., Lambeir, A.-M., Meester, I.D., Man, J.G.D., and Winter, B.Y.D. (2017). Regulation of intestinal permeability: The role of proteases. World J. Gastroenterol. 23, 2106–2123. 10.3748/wjg.v23.i12.2106.

11. Steck, N., Mueller, K., Schemann, M., and Haller, D. (2012). Bacterial proteases in IBD and IBS. Gut 61, 1610. 10.1136/gutjnl-2011-300775.

12. Edogawa, S., Edwinson, A.L., Peters, S.A., Chikkamenahalli, L.L., Sundt, W., Graves, S., Gurunathan, S.V., Breen-Lyles, M., Johnson, S., Dyer, R., et al. (2019). Serine proteases as luminal mediators of intestinal barrier dysfunction and symptom severity in IBS. Gut 69, 62–73. 10.1136/gutjnl-2018-317416.

13. Rangan, K.J., Pedicord, V.A., Wang, Y.-C., Kim, B., Lu, Y., Shaham, S., Mucida, D., and Hang, H.C. (2016). A secreted bacterial peptidoglycan hydrolase enhances tolerance to enteric pathogens. Science 353, 1434–1437. 10.1126/science.aaf3552.

14. Thursby, E., and Juge, N. (2017). Introduction to the human gut microbiota. Biochem. J. 474, 1823 1836. 10.1042/bcj20160510.

15. Heuberger, D.M., and Schuepbach, R.A. (2019). Protease-activated receptors (PARs): mechanisms of action and potential therapeutic modulators in PAR-driven inflammatory diseases. Thrombosis J. 17, 4. 10.1186/s12959-019-0194-8.

16. Maharshak, N., Huh, E.Y., Paiboonrungruang, C., Shanahan, M., Thurlow, L., Herzog, J., Djukic, Z., Orlando, R., Pawlinski, R., Ellermann, M., et al. (2015). Enterococcus faecalis Gelatinase Mediates Intestinal Permeability via Protease-Activated Receptor 2. Infect. Immun. 83, 2762–2770. 10.1128/iai.00425-15.

17. Han, S., Treuren, W.V., Fischer, C.R., Merrill, B.D., DeFelice, B.C., Sanchez, J.M., Higginbottom, S.K., Guthrie, L., Fall, L.A., Dodd, D., et al. (2021). A metabolomics pipeline for the mechanistic interrogation of the gut microbiome. Nature 595, 415–420. 10.1038/s41586-021-03707-9.

18. Caminero, A., Guzman, M., Libertucci, J., and Lomax, A.E. (2023). The emerging roles of bacterial proteases in intestinal diseases. Gut Microbes 15, 2181922. 10.1080/19490976.2023.2181922.

19. Denadai-Souza, A., Bonnart, C., Tapias, N.S., Marcellin, M., Gilmore, B., Alric, L., Bonnet, D., Burlet-Schiltz, O., Hollenberg, M.D., Vergnolle, N., et al. (2018). Functional Proteomic Profiling of Secreted Serine Proteases in Health and Inflammatory Bowel Disease. Sci. Rep. 8, 7834. 10.1038/s41598-018-26282-y.

20. Peach, C.J., Edgington-Mitchell, L.E., Bunnett, N.W., and Schmidt, B.L. (2023). Protease-activated receptors in health and disease. Physiol. Rev. 103, 717–785. 10.1152/physrev.00044.2021.

21. Nguyen, T.T.H., Myrold, D.D., and Mueller, R.S. (2019). Distributions of Extracellular Peptidases Across Prokaryotic Genomes Reflect Phylogeny and Habitat. Front. Microbiol. 10, 413. 10.3389/fmicb.2019.00413.

22. Liu, Y., Patricelli, M.P., and Cravatt, B.F. (1999). Activity-based protein profiling: The serine hydrolases. Proc. Natl. Acad. Sci. 96, 14694–14699. 10.1073/pnas.96.26.14694.

23. Keller, L.J., Babin, B.M., Lakemeyer, M., and Bogyo, M. (2020). Activity-based protein profiling in bacteria: Applications for identification of therapeutic targets and characterization of microbial communities. Curr. Opin. Chem. Biol. 54, 45–53. 10.1016/j.cbpa.2019.10.007.

24. Consortium, T.U. (2022). UniProt: the Universal Protein Knowledgebase in 2023. Nucleic Acids Res. 10.1093/nar/gkac1052.

25. Paysan-Lafosse, T., Blum, M., Chuguransky, S., Grego, T., Pinto, B.L., Salazar, G.A., Bileschi, M.L., Bork, P., Bridge, A., Colwell, L., et al. (2022). InterPro in 2022. Nucleic Acids Res. 10.1093/nar/gkac993.

26. Zou, Y., Xue, W., Luo, G., Deng, Z., Qin, P., Guo, R., Sun, H., Xia, Y., Liang, S., Dai, Y., et al. (2019). 1,520 reference genomes from cultivated human gut bacteria enable functional microbiome analyses. Nat. Biotechnol. 37, 179–185. 10.1038/s41587-018-0008-8.

27. García-Bayona, L., and Comstock, L.E. (2019). Streamlined Genetic Manipulation of Diverse Bacteroides and Parabacteroides Isolates from the Human Gut Microbiota. Mbio 10, e01762–19. 10.1128/mbio.01762-19.

28. Jumper, J., Evans, R., Pritzel, A., Green, T., Figurnov, M., Ronneberger, O., Tunyasuvunakool, K., Bates, R., Žídek, A., Potapenko, A., et al. (2021). Highly accurate protein structure prediction with AlphaFold. Nature 596, 583–589. 10.1038/s41586-021-03819-2.

29. Latorre, R., Hegron, A., Peach, C.J., Teng, S., Tonello, R., Retamal, J.S., Klein-Cloud, R., Bok, D., Jensen, D.D., Gottesman-Katz, L., et al. (2022). Mice expressing fluorescent PAR 2 reveal that endocytosis mediates colonic inflammation and pain. Proc. Natl. Acad. Sci. 119. 10.1073/pnas.2112059119.

30. Co, J.Y., Margalef-Català, M., Li, X., Mah, A.T., Kuo, C.J., Monack, D.M., and Amieva, M.R. (2019). Controlling Epithelial Polarity: A Human Enteroid Model for Host-Pathogen Interactions. Cell Reports 26, 2509–2520.e4. 10.1016/j.celrep.2019.01.108.

31. Vergnolle, N., Bunnett, N.W., Sharkey, K.A., Brussee, V., Compton, S.J., Grady, E.F., Cirino, G., Gerard, N., Basbaum, A.I., Andrade-Gordon, P., et al. (2001). Proteinase-activated receptor-2 and hyperalgesia: A novel pain pathway. Nat. Med. 7, 821–826. 10.1038/89945.

32. Jimenez-Vargas, N.N., Pattison, L.A., Zhao, P., Lieu, T., Latorre, R., Jensen, D.D., Castro, J., Aurelio, L., Le, G.T., Flynn, B., et al. (2018). Protease-activated receptor-2 in endosomes signals persistent pain of irritable bowel syndrome. Proc. Natl. Acad. Sci. 115, E7438–E7447. 10.1073/pnas.1721891115.

33. Cheng, R.K.Y., Fiez-Vandal, C., Schlenker, O., Edman, K., Aggeler, B., Brown, D.G., Brown, G.A., Cooke, R.M., Dumelin, C.E., Doré, A.S., et al. (2017). Structural insight into allosteric modulation of protease-activated receptor 2. Nature 545, 112–115. 10.1038/nature22309.

34. Tonello, R., Anderson, W.B., Davidson, S., Escriou, V., Yang, L., Schmidt, B.L., Imlach, W.L., and Bunnett, N.W. (2023). The contribution of endocytosis to sensitization of nociceptors and synaptic transmission in nociceptive circuits. PAIN 164, 1355–1374. 10.1097/j.pain.0000000000002826.

35. Martini, E., Krug, S.M., Siegmund, B., Neurath, M.F., and Becker, C. (2017). Mend Your Fences The Epithelial Barrier and its Relationship With Mucosal Immunity in Inflammatory Bowel Disease. Cell Mol. Gastroenterol. Hepatol. 4, 33–46. 10.1016/j.jcmgh.2017.03.007.

36. Zheng, D., Liwinski, T., and Elinav, E. (2020). Interaction between microbiota and immunity in health and disease. Cell Res. 30, 492–506. 10.1038/s41422-020-0332-7.

37. Forbes, J.D., Domselaar, G.V., and Bernstein, C.N. (2016). The Gut Microbiota in Immune-Mediated Inflammatory Diseases. Front. Microbiol. 7, 1081. 10.3389/fmicb.2016.01081.

38. Santiago, A., Hann, A., Constante, M., Rahmani, S., Libertucci, J., Jackson, K., Rueda, G., Rossi, L., Ramachandran, R., Ruf, W., et al. (2023). Crohn’s disease proteolytic microbiota enhances inflammation through PAR2 pathway in gnotobiotic mice. Gut Microbes 15, 2205425. 10.1080/19490976.2023.2205425.

39. Carroll, I.M., and Maharshak, N. (2013). Enteric bacterial proteases in inflammatory bowel disease-pathophysiology and clinical implications. World J. Gastroentero. 19, 7531–7543. 10.3748/wjg.v19.i43.7531.

40. Rolland-Fourcade, C., Denadai-Souza, A., Cirillo, C., Lopez, C., Jaramillo, J.O., Desormeaux, C., Cenac, N., Motta, J.-P., Larauche, M., Taché, Y., et al. (2017). Epithelial expression and function of trypsin-3 in irritable bowel syndrome. Gut 66, 1767. 10.1136/gutjnl-2016-312094.

41. Motta, J., Magne, L., Descamps, D., Rolland, C., Squarzoni–Dale, C., Rousset, P., Martin, L., Cenac, N., Balloy, V., Huerre, M., et al. (2011). Modifying the Protease, Antiprotease Pattern by Elafin Overexpression Protects Mice From Colitis. Gastroenterology 140, 1272–1282. 10.1053/j.gastro.2010.12.050.

42. Baker, C.C., Sessenwein, J.L., Wood, H.M., Yu, Y., Tsang, Q., Alward, T.A., Vargas, N.N.J., Omar, A.A., McDonnel, A., Segal, J.P., et al. (2024). Protease-Induced Excitation of Dorsal Root Ganglion Neurons in Response to Acute Perturbation of the Gut Microbiota Is Associated With Visceral and Somatic Hypersensitivity. Cell. Mol. Gastroenterol. Hepatol. 18, 101334. 10.1016/j.jcmgh.2024.03.006.

43. Zhao, P., Metcalf, M., and Bunnett, N.W. (2014). Biased Signaling of Protease-Activated Receptors. Front. Endocrinol. 5, 67. 10.3389/fendo.2014.00067.

44. Cenac, N., Garcia-Villar, R., Ferrier, L., Larauche, M., Vergnolle, N., Bunnett, N.W., Coelho, A.-M., Fioramonti, J., and Bueno, L. (2003). Proteinase-Activated Receptor-2-Induced Colonic Inflammation in Mice: Possible Involvement of Afferent Neurons, Nitric Oxide, and Paracellular Permeability. J. Immunol. 170, 4296–4300. 10.4049/jimmunol.170.8.4296.

45. Jacob, C., Yang, P.-C., Darmoul, D., Amadesi, S., Saito, T., Cottrell, G.S., Coelho, A.-M., Singh, P., Grady, E.F., Perdue, M., et al. (2005). Mast Cell Tryptase Controls Paracellular Permeability of the Intestine ROLE OF PROTEASE-ACTIVATED RECEPTOR 2 AND β-ARRESTINS*. J. Biol. Chem. 280, 31936–31948. 10.1074/jbc.m506338200.

46. Cattaruzza, F., Lyo, V., Jones, E., Pham, D., Hawkins, J., Kirkwood, K., Valdez-Morales, E., Ibeakanma, C., Vanner, S.J., Bogyo, M., et al. (2011). Cathepsin S Is Activated During Colitis and Causes Visceral Hyperalgesia by a PAR2-Dependent Mechanism in Mice. Gastroenterology 141, 1864–1874.e3. 10.1053/j.gastro.2011.07.035.

47. Róka, R., Demaude, J., Cenac, N., Ferrier, L., Salvador-cartier, C., Garcia-villar, R., Fioramonti, J., and Bueno, L. (2007). Colonic luminal proteases activate colonocyte proteinase-activated receptor-2 and regulate paracellular permeability in mice. Neurogastroenterol. Motil. 19, 57–65. 10.1111/j.1365-2982.2006.00851.x.

48. Rondeau, L.E., Luz, B.B.D., Santiago, A., Bermudez-Brito, M., Hann, A., Palma, G.D., Jury, J., Wang, X., Verdu, E.F., Galipeau, H.J., et al. (2024). Proteolytic bacteria expansion during colitis amplifies inflammation through cleavage of the external domain of PAR2. Gut Microbes 16, 2387857. 10.1080/19490976.2024.2387857.

49. Gibson, S.A.W., and Macfarlane, G.T. (1988). Characterization of Proteases Formed by Bacteroides fragilis. Microbiology+ 134, 2231–2240. 10.1099/00221287-134-8-2231.

50. Riepe, S.P., Goldstein, J., and Alpers, D.H. (1980). Effect of Secreted Bacteroides Proteases on Human Intestinal Brush Border Hydrolases. J. Clin. Invest. 66, 314–322. 10.1172/jci109859.

51. Macfarlane, G.T., Macfarlane, S., and Gibson, G.R. (1992). Synthesis and release of proteases by Bacteroides fragilis. Curr. Microbiol. 24, 55–59. 10.1007/bf01570100.

52. Wexler, H.M. (2007). Bacteroides: the Good, the Bad, and the Nitty-Gritty. Clin. Microbiol. Rev. 20, 593–621. 10.1128/cmr.00008-07.

53. Zafar, H., and Saier, M.H. (2021). Gut Bacteroides species in health and disease. Gut Microbes 13, 1848158. 10.1080/19490976.2020.1848158.

54. Xu, J.H., Jiang, Z., Solania, A., Chatterjee, S., Suzuki, B., Lietz, C.B., Hook, V.Y.H., O’Donoghue, A.J., and Wolan, D.W. (2018). A Commensal Dipeptidyl Aminopeptidase with Specificity for N-terminal Glycine Degrades Human-produced Antimicrobial Peptides In Vitro. ACS Chem. Biol. 13. 10.1021/acschembio.8b00420.

55. Roncase, E.J., González-Páez, G.E., and Wolan, D.W. (2019). X-ray Structures of Two Bacteroides thetaiotaomicron C11 Proteases in Complex with Peptide-Based Inhibitors. Biochemistry 58, 1728–1737. 10.1021/acs.biochem.9b00098.

56. Keller, L.J., Nguyen, T.H., Liu, L.J., Hurysz, B.M., Lakemeyer, M., Guerra, M., Gelsinger, D.J., Chanin, R., Ngo, N., Lum, K.M., et al. (2023). Chemoproteomic identification of a DPP4 homolog in Bacteroides thetaiotaomicron. Nat. Chem. Biol. 19, 1469–1479. 10.1038/s41589-023-01357-8.

57. Roncase, E.J., Moon, C., Chatterjee, S., González-Páez, G.E., Craik, C.S., O’Donoghue, A.J., and Wolan, D.W. (2017). Substrate Profiling and High Resolution Co-complex Crystal Structure of a Secreted C11 Protease Conserved across Commensal Bacteria. ACS Chem. Biol. 12, 1556–1565. 10.1021/acschembio.7b00143.

58. Pierce, J.V., Fellows, J.D., Anderson, D.E., and Bernstein, H.D. (2021). A clostripain-like protease plays a major role in generating the secretome of enterotoxigenic Bacteroides fragilis. Mol. Microbiol. 115, 290–304. 10.1111/mmi.14616.

59. Choi, V.M., Herrou, J., Hecht, A.L., Teoh, W.P., Turner, J.R., Crosson, S., and Wardenburg, J.B. (2016). Activation of Bacteroides fragilis toxin by a novel bacterial protease contributes to anaerobic sepsis. Nat. Med. 22, 563–567. 10.1038/nm.4077.

60. Galipeau, H.J., Caminero, A., Turpin, W., Bermudez-Brito, M., Santiago, A., Libertucci, J., Constante, M., Garay, J.A.R., Rueda, G., Armstrong, S., et al. (2021). Novel Fecal Biomarkers That Precede Clinical Diagnosis of Ulcerative Colitis. Gastroenterology 160, 1532–1545. 10.1053/j.gastro.2020.12.004.

61. Nomura, K., Ishikawa, D., Okahara, K., Ito, S., Haga, K., Takahashi, M., Arakawa, A., Shibuya, T., Osada, T., Kuwahara-Arai, K., et al. (2021). Bacteroidetes Species Are Correlated with Disease Activity in Ulcerative Colitis. J. Clin. Medicine 10, 1749. 10.3390/jcm10081749.

62. Porras, A.M., Zhou, H., Shi, Q., Xiao, X., Bank, J.L.C., Longman, R., and Brito, I.L. (2022). Inflammatory Bowel Disease-Associated Gut Commensals Degrade Components of the Extracellular Matrix. mBio 13, e02201–22. 10.1128/mbio.02201-22.

63. Mills, R.H., Dulai, P.S., Vázquez-Baeza, Y., Sauceda, C., Daniel, N., Gerner, R.R., Batachari, L.E., Malfavon, M., Zhu, Q., Weldon, K., et al. (2022). Multi-omics analyses of the ulcerative colitis gut microbiome link Bacteroides vulgatus proteases with disease severity. Nat. Microbiol. 7, 262–276. 10.1038/s41564-021-01050-3.

64. Patrick, S. (2022). A tale of two habitats: Bacteroides fragilis, a lethal pathogen and resident in the human gastrointestinal microbiome. Microbiology 168. 10.1099/mic.0.001156.

65. Sofi, M.H., Wu, Y., Ticer, T., Schutt, S., Bastian, D., Choi, H.-J., Tian, L., Mealer, C., Liu, C., Westwater, C., et al. (2021). A single strain of Bacteroides fragilis protects gut integrity and reduces GVHD. JCI Insight 6, e136841. 10.1172/jci.insight.136841.

66. He, Q., Niu, M., Bi, J., Du, N., Liu, S., Yang, K., Li, H., Yao, J., Du, Y., and Duan, Y. (2023). Protective effects of a new generation of probiotic Bacteroides fragilis against colitis in vivo and in vitro. Sci. Rep. 13, 15842. 10.1038/s41598-023-42481-8.

67. Lee, Y.K., Mehrabian, P., Boyajian, S., Wu, W.-L., Selicha, J., Vonderfecht, S., and Mazmanian, S.K. (2018). The Protective Role of Bacteroides fragilis in a Murine Model of Colitis-Associated Colorectal Cancer. MSphere 3, e00587–18. 10.1128/msphere.00587-18.

68. Swidsinski, A., Weber, J., Loening-Baucke, V., Hale, L.P., and Lochs, H. (2005). Spatial Organization and Composition of the Mucosal Flora in Patients with Inflammatory Bowel Disease. J. Clin. Microbiol. 43, 3380–3389. 10.1128/jcm.43.7.3380-3389.2005.

69. Kumbhari, A., Cheng, T.N.H., Ananthakrishnan, A.N., Kochar, B., Burke, K.E., Shannon, K., Lau, H., Xavier, R.J., and Smillie, C.S. (2024). Discovery of disease-adapted bacterial lineages in inflammatory bowel diseases. Cell Host Microbe 32, 1147–1162.e12. 10.1016/j.chom.2024.05.022.

70. Becker, H.E.F., Jamin, C., Bervoets, L., Boleij, A., Xu, P., Pierik, M.J., Stassen, F.R.M., Savelkoul, P.H.M., Penders, J., and Jonkers, D.M.A.E. (2021). Higher Prevalence of Bacteroides fragilis in Crohn’s Disease Exacerbations and Strain-Dependent Increase of Epithelial Resistance. Front. Microbiol. 12, 598232. 10.3389/fmicb.2021.598232.

71. Ning, L., Zhou, Y.-L., Sun, H., Zhang, Y., Shen, C., Wang, Z., Xuan, B., Zhao, Y., Ma, Y., Yan, Y., et al. (2023). Microbiome and metabolome features in inflammatory bowel disease via multi-omics integration analyses across cohorts. Nat. Commun. 14, 7135. 10.1038/s41467-023-42788-0.

72. Amadesi, S., Cottrell, G.S., Divino, L., Chapman, K., Grady, E.F., Bautista, F., Karanjia, R., Barajas-Lopez, C., Vanner, S., Vergnolle, N., et al. (2006). Protease-activated receptor 2 sensitizes TRPV1 by protein kinase Cepsilon- and A-dependent mechanisms in rats and mice. J. Physiol. 575, 555–571. 10.1113/jphysiol.2006.111534.

73. Barbara, G., Stanghellini, V., Giorgio, R.D., Cremon, C., Cottrell, G.S., Santini, D., Pasquinelli, G., Morselli-Labate, A.M., Grady, E.F., Bunnett, N.W., et al. (2004). Activated mast cells in proximity to colonic nerves correlate with abdominal pain in irritable bowel syndrome. Gastroenterology 126, 693–702. 10.1053/j.gastro.2003.11.055.

74. Steinhoff, M., Vergnolle, N., Young, S.H., Tognetto, M., Amadesi, S., Ennes, H.S., Trevisani, M., Hollenberg, M.D., Wallace, J.L., Caughey, G.H., et al. (2000). Agonists of proteinase-activated receptor 2 induce inflammation by a neurogenic mechanism. Nat. Med. 6, 151–158. 10.1038/72247.

