## Supplementary material for "Identification of a secreted protease from *Bacteroides fragilis* that induces intestinal pain and inflammation by cleavage of PAR_2_": Materials and Methods

**Generation of His-PAR_2_-NTD-Cy5 peptide by expressed protein ligation.**

Design: The PAR_2_-NTD-peptide, consisting of the 46 amino acids spanning human PAR_2_-NTD and a N-terminal hexahistidine (His_6_)-sequence, was fused to the intein Mxe GyrA. To improve efficiency of native chemical ligation and protein quantification, a two amino acid linker sequence (WG) was introduced before the GyrA sequence. Nucleotide and protein sequences can be found in **Supplementary Table S5**.

Recombinant protein expression and purification of His-PAR_2_NTD-GyrA: Plasmid pET28a_His-PAR_2_NTD-GyrA was transformed into chemically competent *E. coli* Rosetta2(DE3) cells with selection using 50 µg ml^−1^ kanamycin and 34 µg ml^−1^ chloramphenicol for expression. For recombinant protein expression, an overnight culture of transformed cells was inoculated into 1 L of LB supplemented with 50 µg ml^−1^ kanamycin and 34 µg ml^−1^ chloramphenicol and grown with shaking until the optical density at 600 nm (OD_600_) = 0.6. After the culture was cold-shocked on ice for 20 min, 0.5 mM isopropyl β-D-1-thiogalactopyranoside (IPTG) was added to the culture to induce protein expression, and the culture was incubated for 20 h at 18 °C. The bacterial pellet was flash-frozen in liquid nitrogen and stored at −80 °C. The pellet was lysed in 30 ml of lysis buffer (50 mM *N*-2-hydroxyethylpiperazine-*N*-2-ethane sulfonic acid (HEPES), 10 mM imidazole, 150 mM NaCl, pH 8) supplemented with 750 mM trehalose by probe sonication on ice (3 min at 30% power in 1 s bursts; 1 min at 60% power in 1 s bursts; 3 min at 30% power in 1 s bursts and 1 min at 60% power in 1 s bursts). Cell debris was pelleted at 38,400 x *g* for 20 min at 4 °C, and the lysate was transferred to a new tube and centrifuged at 12,000 x *g* for 15 min at 4 °C. The lysate was then clarified sequentially through a 5-µm and a 1-µm filter before purification. The clarified lysate was injected into an ÄKTA purifier FPLC system (GE Healthcare) and separated with a 5 mL HisTrap FF crude column (GE Healthcare). The column was subsequently washed with lysis buffer supplemented with 10 mM imidazole, 20 mM imidazole, and 40 mM imidazole. The protein was eluted with lysis buffer supplemented with 300 mM imidazole, and fractions containing protein (as determined by the ultraviolet (UV) trace and SDS–PAGE) were combined. The imidazole-concentration of the final protein sample was reduced to ca. 30 mM by consecutive buffer-exchange with buffer (200 mM HEPES, 150 mM NaCl, pH 8).

Native chemical ligation: On-resin native chemical ligation was used to C-terminally conjugate the His_6_-PAR_2_-peptide with Cys-PEG_4_-Cy5 (see **Supplementary Note 1**). To do so, 1.2 mL of 300 µM His_6_-PAR_2_-GyrA (360 nmol) in thioesterification buffer (200 mM HEPS, 150 mM NaCl, pH 8) was added to 2.4 mL of 150 mM sodium 2-mercaptoethanesulfonate (Mesna) in freshly degassed thioesterification buffer and incubated at 4°C for 18 h while gently rocking. The sample was diluted with 1.8 mL of pulldown-buffer (200 mM HEPES, 20 mM NaCl, 20 mM imidazole, pH 7.2) and 300 µL of pre-washed Ni-NTA agarose beads (Qiagen) was added. Pulldown of His_6_-PAR_2_-Mesna was performed at 4°C for 2 h while gently rocking. The supernatant containing cleaved GyrA was removed, and the beads were washed six times with 500 µL of pulldown-buffer. For native chemical ligation, beads were resuspended in freshly degassed NCL-Buffer (200 mM HEPES, 20 mM NaCl, 20 mM Tris(2-carboxyethyl)phosphine (TCEP), 100 mM 4-Mercaptophenylacetic acid (MPAA), pH 7.4), 18 µL of 50 mM Cys-PEG_4_-Cy5 (2.5 eq. based on original His_6_-PAR_2_-Mesna) was added. The reaction was incubated under an argon atmosphere at 4 °C for 16 h while gently rocking. The supernatant was removed and beads were washed six times with 300 µL of wash-buffer-1 (200 mM HEPES, 20 mM NaCl, 1 mM TCEP, pH 7.2). For alkylation, beads were resuspended in 300 µL of freshly made alkylation-buffer (200 mM HEPES, 20 mM NaCl, 1 mM TCEP, 5 µM iodoacetamide) and incubated at rt in the dark for 90 min. The supernatant was removed and beads were washed three times with 500 µL of wash-buffer-2 (200 mM HEPES, 20 mM NaCl, pH = 7.2) and three times with 500 µL of ultrapure water. His_6_-PAR_2_NTD-Cy5 peptide was eluted from the Ni-NTA beads by treatment with four times 150 µL of 0.5% TFA in ultrapure water. The peptide was purified by preparative HPLC (column: Aeris 3.6 µm Widepore C4 200, 250 x 4.6 mm; solvents: A: 0.1% TFA in water, B: 0.1% TFA in MeCN; gradient: 5% B to 80% B in 30 min) and lyophilized. The final peptide-TFA salt (1.13 mg; 75 nmol, 21% based on Cy5-absorbance) was dissolved in 0.05% formic acid in ultrapure water and its concentration was determined by Cy5 absorbance (e = 271,000 M^-1^cm^-1^ at 649 nm).

**Bacterial culture media and growth conditions.** Liquid handling and cultivations for all strains (except for *E. coli* strains for molecular cloning and recombinant protein expression) were performed inside an anaerobic chamber (Coy Lab Products) with an atmosphere of 5% H_2_, 5% CO_2,_ and 90% N_2_. Unless stated otherwise, bacteria were grown in Mega Medium (see **Supplementary Note 2**).^1^ Identity of the strains was verified by 16S rRNA sequencing, as previously described.^1^

**PAR_2_-proteolysis screening.**

Cultivation and supernatant generation: Microbes were cultivated in 96-well plates with every well corresponding to an individual bacterial strain, as previously described.^1^ Starter cultures (1 mL scale) were grown for 24 h at 37 °C without shaking. Growth was verified by OD_600_ measurement. For main cultures, Mega Medium (1 mL) was inoculated from overnight cultures (1:50) and cultivated for 24 h without shaking. Growth was monitored in parallel in a 96-well plate format via OD_600_-measurements using a Cerillo® Stratus plate-reader mounted on a plate shaker (200 rpm). Culture plates were taken outside the anaerobic chamber, centrifuged (4500 x g, 4°C, 15 min) and supernatants were sterile filtered using 0.22 µm filter plates. The cell-free supernatants were aliquoted, snap-frozen and stored at -80 °C until used for activity assays.

Proteolysis and affinity-pulldown: Supernatants (9 µL) and His-PAR_2_-NTD-Cy5 substrate (1 µL of 20 µM in 0.01% FA) were incubated for 3 h at 37 °C. Next, samples were cooled in a cold-block and centrifuged. For affinity-pulldown, Ni-NTA magnetic beads (PureCube®) were washed in pulldown-buffer (200 mM HEPES, 0.025 % Tween-80, pH = 8.8) and 1 µL of original bead suspension in 16 µL of pulldown-buffer were added to each sample. Pulldown was performed for 2 h at 4 °C while shaking. Samples were centrifuged, beads were sedimented using a magnetic rack and 10 µL of the supernatant was transferred to a small-volume 384 well-plate (Greiner). Fluorescence of the proteolytically cleaved peptide fragments was determined using a Cytation3 plate-reader (excitation: 640 nm, emission: 670 nm). For each sample, PAR_2_-processing activity was normalized to trypsin (100% activity) and Mega Medium-background (0% activity). The screening results were plotted onto a phylogenetic tree (based on 16S rRNA sequences) of the strain tested via iTol (https://itol.embl.de).

**Validation of PAR_2_-processing activity and inhibition by covalent inhibitors.** To validate the PAR_2_-processing activity, verified strains (via 16S rRNA sequencing) with more than 50% PAR_2_-processing activity in the initial screen were cultivated individually. To do so, 10 mL Mega Medium was inoculated with 0.1 mL of an overnight-culture and cultivated at 37 °C without shaking. Growth was monitored in parallel in a 96-well plate format via OD_600_-measurements using a Cerillo® Stratus plate-reader mounted on a plate shaker (200 rpm). When a culture reached early stationary phase, the culture tube was centrifuged (4500 x g, 4°C, 15 min) and supernatants were sterile filtered using a 0.22 µm PES syringe filter. Supernatants were then subjected to activity-assays. For each assay, 9 µL of each supernatant was pre-incubated with 0.2 µL DMSO (control) or covalent inhibitor (50x stock concentration) at 37 °C for 1h. Samples were centrifuged and 1 µL of His-PAR_2_-NTD-Cy5 substrate (20 µM in 0.01% FA) was added to each well. Proteolysis was performed for 1.5 h (instead of 3 h for the initial screen) at 37°C. Affinity-pulldown and fluorescence measurements were performed as described above. For each sample, PAR_2_-processing activity was normalized to trypsin (100% activity) and Mega Medium-background (0% activity). Experiments were performed in duplicates (n = 2).

**Gel-based ABPP.** *B. fragilis NCTC 9343* culture supernatants were harvested at early stationary phase (ca. 9h cultivation) via centrifugation (5000 x g, 15 min, 4 °C) and sterile-filtration (0.22 µm PES sterile filter). Supernatants were then concentrated 5-fold using 10 kDa MW-cutoff-filters. For labeling, 60 µL of 5-fold supernatants were incubated with 0.6 µL of DMSO or MLP7 (0.5 mM, 2 mM, or 10 mM) for 1 h at 37 °C. Subsequently, samples were treated with 0.6 µL of DMSO or FP-TMR (0.1 mM or 1 mM) for 1 h at 37 °C. Samples were precipitated by chloroform-methanol precipitation. In brief, each sample was treated with methanol (240 µL), chloroform (60 µL), and water (180 µL) with mixing after each addition step. After centrifugation (18000 x g, 5 min, rt) the upper layer was carefully removed. Methanol (180 µL) was added and the sample was mixed and centrifuged (18000 x g, 5 min, rt) again. The full supernatant was removed and the remaining pellet dissolved in 1x Laemmli-Buffer. Labeled proteins were separated via SDS–PAGE. In-gel fluorescence was visualized using the Cy3 channel on Typhoon 9410 Imager (Amersham Biosciences).

**MS-based ABPP sample preparation and analysis.** *B. fragilis NCTC 9343* cultures were grown in 110 mL of Mega Medium in triplicates for 9 h until early-stationary phase. Cultures were centrifuged (5000 x g, 15 min, 4 °C) to pellet bacterial cells and supernatants were filtered using a 0.22 µm PES sterile filter. 100 mL of the supernatants were concentrated to a final volume of 20 mL using a 10 kDa MW-cutoff concentrator. For each biological replicate, the supernatant was aliquoted into three separate tubes (6 mL each), representing the three labeling conditions “DMSO”, “Probe” and “Competition”. Supernatants were incubated with 60 µL DMSO (for “DMSO” and “Probe”) or 60 µL of 10 mM MLP7 (for “Competition”) at 37 °C for 1 h. Subsequently, samples were treated with 60 µL of DMSO (for “DMSO”) or 60 µL of 1 mM FP-Biotin (for “Probe” and “Competition”) and incubated at 37 °C for 1 h. For precipitation, 606 µL of 100% trichloroacetic acid (TCA) solution was added to each sample, incubated on ice for 30 min, and centrifuged (12,000 x g, 4 °C for 15 min). The supernatants were removed, the pellets were resuspended in a mix of 0.5 mL PBS and 4.5 mL acetone and centrifuged (12,000 x g, 4 °C for 15 min). The supernatants were removed, and the pellets washed again with a mix of 0.375 mL PBS and 3.4 mL acetone. Protein pellets were solubilized in 0.5 ml PBS containing 0.4% (w/v) SDS and transferred to Lo-bind Eppendorf tubes containing 50 μL of avidin-agarose bead slurry (Sigma Aldrich) pre-equilibrated with 0.4% SDS (w/v) in PBS (3 × 1 mL, 400 x *g*, 5 min, rt), and rotated for 1 h at room temperature. The beads were then washed with 0.4% (w/v) SDS in PBS (3 × 1 mL), 6 M urea (2 × 1 mL) and finally PBS (3 × 1 mL). Beads were resuspended in 200 μL X-buffer (7 M urea, 2 M thiourea in 20mM HEPES buffer, pH 7.5). Upon reduction with 5 mM TCEP (2 μL of 500 mM stock in ddH_2_O) for 1 h at 37°C, proteins were alkylated using 10 mM IAA (4 μL of 500 mM stock in ddH_2_O) for 30 min at 25°C and samples were quenched with 10 mM DTT (4 μL of 500 mM stock in ddH_2_O) for 30 min at 25°C. Enzymatic digestion using LysC (1 μL of 0.5 μg/μL, Wako, MS-grade) was first carried out for 2 h at 25°C, upon which samples were diluted with triethylammonium bicarbonate (TEAB) buffer (600 μL of 50 mM stock in ddH_2_O) and digested with trypsin (1.5 μL of 0.5 μg/μL in 50 mM acetic acid, Promega, sequencing grade) for a further 16 h at 37°C. Samples were acidified to 1% (v/v) FA and desalted using SepPak® C18 cartridges (50 mg, Waters) with a vacuum manifold. The cartridges were first washed with ACN (2 × 1 ml) and equilibrated with 0.1% (v/v) TFA (3 × 1 mL) prior to loading the samples. After washing with 0.1% (v/v) TFA (3 × 1 mL) and 0.5% (v/v) FA (1 × 0.5 mL), peptides were eluted in 80% (v/v) ACN containing 0.5% FA (3 × 0.25 mL) and freeze-dried using a speedvac centrifuge. For tandem-mass-tag (TMT)-labeling, samples were redissolved in 23 µL of 100 mM HEPES-Buffer (pH = 8.5) and treated with 7.5 µL of respective TMT-reagent (Thermo Scientific 10-plex kit, each label was dissolved to 13 µg/µL in anhydrous acetonitrile) and incubated (25 °C, 500 rpm) for 1 h. The reactions were stopped by adding 0.3 µL of 50% hydroxylamine and incubating it (25 °C, 500 rpm) for 15 min. Samples were freeze-dried using a speedvac centrifuge, desalted using C18-tips (Pierce® C18 tips, 100 µL, Thermo Scientific) according to the manufacturer’s protocol and freeze-dried using a speedvac centrifuge. For measurement, samples were dissolved in 20 µL of 1% formic acid in water. MS analysis was performed using an Orbitrap Eclipse Tribrid mass spectrometer (Thermo Scientific, San Jose, CA, USA) with liquid chromatography using an Acquity M-Class UPLC (Waters Corporation, Milford, MA, USA). A flow rate of 300 nL/min was used, where mobile phase A was 0.2% formic acid in water and mobile phase B was 0.2% formic acid in acetonitrile. Analytical columns were prepared in-house with an I.D. of 100 microns pulled to a nanospray emitter using a P2000 laser puller (Sutter Instrument, Novato, CA, USA). The column was packed using C18 reprosil Pur 1.8-micron stationary phase (Dr. Maisch) to a length of ~25 cm. Peptides were directly injected onto the analytical column using a gradient (2%–45% B, followed by a high-B wash) of 180 min. MS data were acquired using an MS3 data-dependent acquisition method. MS1 profile scans were acquired in the Orbitrap (resolution, 120,000; scan range, 400–1600 *m/z* ; AGC target, 4 × 10^5^ and maximum injection time, 50 ms). Monoisotopic peak determination was set to ‘peptide’. Only charge states 2 – 6 were included. Dynamic exclusion was enabled (repeat count, *n* = 1; exclusion duration, 30 s; mass tolerance: low, 10 ppm and high, 10 ppm, excluding isotopes). An intensity threshold of 5 × 10^3^ was set. Data-dependent MS2 spectra were acquired in centroid mode across a mass range of 400–1,600 *m/z*. Precursor ions were isolated using the quadrupole (isolation window, 0.7 *m/z*), fragmented using collision-induced dissociation (CID) (collision energy, 35%; activation time, 10 ms and activation Q, 0.25) and detected in the ion trap (scan range mode, auto *m/z* normal; scan rate, turbo; AGC target, 1 × 10^4^ and maximum injection time, 35 ms). Data-dependent MS3 spectra were acquired in the Orbitrap (resolution, 50,000; scan range, 100–500 *m/z*; AGC target, 1 × 10^5^and maximum injection time, 200 ms) following higher-energy collisional dissociation (HCD) activation (collision energy, 55%) using Synchronous Precursor Selection from up to 10 precursors. Peptide and protein identifications were performed using MaxQuant (version 2.3.0.3)^2^ with Andromeda as the search engine. Group-specific parameters were set to ‘Reporter ion MS3’ with 10plex TMT isobaric labels for N-terminal and lysine residue modification selected. Reporter mass tolerance was set to 0.003 Da. The following parameters were used: carbamidomethylation of cysteines as fixed modifications, oxidation of methionine and acetylation of N-terminus as dynamic modifications, and trypsin/P as the proteolytic enzyme. Default settings were used for all other parameters. Searches were performed against the UniProt database for *B. fragilis* NCTC 9343 (proteome ID: UP000006731, downloaded on 11.02.2022.) with the following parameters: minimum peptide length, 6; maximum peptide mass, 6,000 Da; minimum peptide length for unspecific search, 6 and maximum peptide length for unspecific search, 40. Identification was performed with at least two unique peptides and quantification only with unique peptides. Statistical analyses were performed with Perseus v.2.0.3.0.^3^ Putative contaminants, reverse hits and proteins identified by side only were removed. Label-free quantitation intensities were log_2_-transformed. Missing values were imputed using a normal distribution (width, 0.3; down-shift, 1.8). *P* values were calculated using a two-sided, two-sample *t*-test. Data are available under: <https://www.ebi.ac.uk/pride/> (Project accession: PXD059166, Token: ceCR6vYkusLh).

**BLAST-N analysis.** BLAST-N analysis (Version BLASTN 2.16.0+ ) against 1,520 reference genomes from cultivated human gut bacteria (Bioproject ID 482748)^4^ was performed at <https://blast.ncbi.nlm.nih.gov/Blast.cgi>^5^ using the nucleotide sequences of Bfp1 (BF9343_2070) and TspA (BF9343_0342) as input with the following parameters: Program selection: discontiguous megablast, Max. target sequences: 500. All other parameters were set to default. The phylogenetic tree for the Blast results was generated with phyloT software (https://phylot.biobyte.de), using the full genomes of 216 species in which the full 1520-member database was clustered, and plotted via iTol (https://itol.embl.de).

**Clean genetic deletion of the bfp1 and tspA genes.** The two clean deletion strains *B. fragilis* NCTC 9343 ∆*bfp1* and ∆*tspA* were generated via two-step allelic exchange based on the vector pLGB30 according to the protocol from Garcia-Bayona et al.^6^ In brief, the plasmids pLGB30-bfp1 and pLGB30-tspA were generated by PCR according to **Supplementary Table S6**. Each plasmid was conjugated in *B. fragilis* NCTC 9343 using *E. coli* S17-1 λ*pir* as the conjugative donor strain. Exconjugants with chromosomally integrated plasmids were recovered on BHIS plates containing 200 µg/mL gentamycin and 6 µg/mL tetracycline. Second crossover events were selected using BHIS plates containing 10 mM L-rhamnose. Deletion of the target genes was confirmed by PCR using suitable primers (see **Supplementary Table S7**).

**Human intestinal organoids.** Human organoid line HC921 established in the Calvin Kuo lab (Stanford University) from healthy ileal tissue was cultured in IntestiCult Organoid Growth Media (Stemcell) supplemented with 10 µM Y27632 (MedChem Express, HY-10583) and 2.5 μM CHIR 99021 (Tocris, 102875-390). TrypLE Express Enzyme (Thermo Fisher Scientific, 12604013) was used to passage organoids before embedding the cells in Matrigel (Corning, 354234) and growing organoids in a 24-well plate with 400 µl of growth medium usually with 1:5 to 1:8 ratio every 7-10 days.

**Polarity reversal and FITC-dextran assay.** To achieve polarity reversal and obtain apical-out organoids, we followed a protocol by Co et al.^7^ and confirmed the polarity reversal using Phalloidin-647 (Cell Signaling Technologies, 8940S). Briefly, organoids 5-7 days after passage were harvested and Matrigel was removed using 5 mM EDTA, 4 °C, 30 minutes. Organoids were washed with AdMEM/F-12 (Thermo Fisher Scientific, 12634010) and then incubated in IntestiCult growth medium for 2 days in ultra-low attachment plates (Corning, 3473) pre-treated with Anti-Adherence Rinsing Solution (Stemcell Technologies, 07010).

Flipped organoids were transferred into 50 µl of phenol-red free DMEM/F-12 (Thermo Fisher Scientific, 21041025) and 10 µl of concentrated bacterial supernatants from WT *B. fragilis*, *B. fragilis* *∆bfp1*-knockout, Mega Medium or 5 mM EDTA were added. Organoids were incubated for 30 minutes at 37 °C before adding 40 µl of 2 mg/ml FITC-dextran (Sigma-Aldrich, 46944-100MG-F) and incubated for 15 minutes. Organoids were then centrifuged for 2 min, 250xg, the supernatant was removed, and organoids were resuspended in the remaining liquid. 5 µl of the sample was transferred onto a microscopy slide and a chamber was created using vacuum grease and a coverslip to prevent mechanical stress. Organoids were then imaged on confocal microscopy. Samples were processed sequentially to ensure precise timing. MFIs were calculated using Zen software (Zeiss). MFI for each organoid was normalized to background for the particular image and values were plotted using GraphPad Prism 10.

**Organoid-derived monolayers and transepithelial resistance measurement.** Organoid-derived monolayers were prepared according to Stemcell’s protocol with minor alterations.^8^  Briefly, Transwells (Corning, 3413) were pre-coated with Matrigel. Organoids were harvested with TrypLE and then trypsin to ensure dissociation into single cells. Cells were seeded at 75,000 cells/well in 100 µl of the recommended medium. Once monolayers were formed (3-4 days), cells were treated apically with the following solutions: 50 µl phenol-red free DMEM/F-12, 10 µl 40x concentrated bacterial supernatants or Mega Medium (final 4x concentrated), PAR_2_-agonistic peptide 2-furoyl-LIGRLO-amide in Mega Medium (final 1 µM), or purified Trypsin in Mega Medium (final 4 nM) and 40 µl of FITC-dextran (2mg/ml). At selected time points, trans-epithelial resistance was measured using EVOM2. Measured values were normalized by subtracting the background (coated well without cells treated apically with Mega Medium) and calculating the resistance per cm^2^.

**Cell lines.** Human embryonic kidney (HEK) 293 cells, which endogenously express PAR_2_, were maintained in culture medium (DMEM + 10% fetal bovine serum (FBS) + penicillin/streptomycin, 50 IU/ml) at 37°C in 5% CO_2_. To make a cell line expressing fluorescently tagged PAR_2_, cDNA encoding human PAR_2_ with N-terminal HA tag and C-terminal mApple fluorescent tag was designed and purchased from Twist Bioscience. HEK293 cells were transfected with 1 µg of the pTwist_CMV_Hygro-HA-PAR_2_-mApple plasmid using PEI (Invitrogen) with a DNA:PEI ratio of 1:6 and cultured for 48 h. Single cells were suspended, plated in 96 well plates, and HA-PAR_2_-mApple expressing cells were selected in culture medium + hygromycin (200 µg/ml). mApple fluorescence was confirmed by microscopy. A single colony with high and uniform expression of HA-PAR_2_-mApple was selected, expanded for further experimentation, and maintained in culture medium with hygromycin (100 µg/ml). HEK293T cells in which PAR_2_ was deleted via CRISPR/Cas genome editing (HEK-PAR_2_-KO) have been described previously.^9^

**Cellular PAR_2_ cleavage assays.** HEK-HA-PAR_2_-mApple cells (30,000 cells) were plated on poly-D-lysine-coated 12 mm round glass coverslips in a 24 well plate and incubated overnight. Cells were washed and incubated in HBSS-H for 30 min. Cells were untreated or incubated with Mega Medium (10x diluted, control), trypsin (10 nM, positive control), WT *B. fragilis* or *B. fragilis* *∆bfp1* supernatant (10x diluted) for 30 min at 37°C. Cells were washed 1x in ice cold HBSS-H, fixed in 4% paraformaldehyde on ice for 20 min and washed with PBS. Cells were incubated in blocking buffer (PBS + 3% normal horse serum + 0.3% saponin, pH 7.4) for 1 h at room temperature (RT). Cells were incubated with the rat anti-HA (1:1000) in PBS at 4°C overnight. Cells were washed 3x in PBS and incubated with donkey anti-rat-AlexaFlour488 (1:1000, ThermoFisher) for 1 h at RT. Cells were washed, incubated with DAPI (1 µM, 5 min), washed, and mounted with ProLong Glass.

**Microscopy.** Cells were imaged on an inverted Leica SP8 confocal microscope or a Leica DMi8 widefield microscope with a 63x objective (1.4 NA). Images were processed with ImageJ (NIH) and figures were made with Adobe Illustrator.

**Animals.** The following strains of mice were used: C57BL/6J wild-type mice (#000664 JAX®); knock-in mice expressing PAR_2_ fused to monomeric ultrastable GFP (*Par_2_-mugfp*); *Par_2_^-/-^* global knockout mice (#0004993 JAX®); mice with PAR_2_ deleted in Nav-1.8+ve nociceptors (*Par_2_^-/-^-Nav1.8)* and PAR_2_-Cre control mice (*Par_2_^+/+^-Cre*)*.* Male mice (8-10 weeks) were used. Mice were maintained in a light-controlled (12-h light/dark cycle) and temperature-controlled (22 ± 4 °C) environment with *ad libitum* access to food and water. The Animal Ethics Committees of New York University and Queen’s University approved animal experiments.

**Patch clamp electrophysiology.** Dorsal root ganglia (DRG) (L1-L5) from C57BL/6J wild-type mice were incubated in collagenase (1 mg/ml, Sigma-Aldrich) and dispase (1 mg/ml, Sigma-Aldrich) in minimal essential medium (Invitrogen) with 10% fetal bovine serum (FBS) for 30 min at 37℃ in a humidified atmosphere (95% air, 5% CO_2_). Neurons were mechanically dispersed by trituration through a fire-polished Pasteur pipette. Neurons were resuspended in neurobasal plus medium with B27 plus supplement (Invitrogen), 10% FBS and penicillin-streptomycin. Neurons were plated onto poly-D-lysine (0.05 mg/ml, Invitrogen) coated glass coverslips and maintained at 37℃ in a humidified atmosphere (95% air, 5% CO_2_). The culture medium was replaced with serum-free neurobasal plus medium 2 h after plating. Patch clamp recordings were made from small diameter (≤ 25 μm) DRG neurons within 30 h of plating. Perforated patch clamp recordings were made using a pipette solution containing amphotericin B (240 μg/ml, Thermo Scientific) in current clamp mode at room temperature (RT). Recordings were made using an amplifier (Axopatch 200B, Molecular Devices) and an analog-digital converter (Digidata 1440A, Molecular Devices) controlled with a PC running pCLAMP 10 software. After a giga-ohm seal was established, neurons were equilibrated for 15 min to achieve membrane perforation and electrical access. Changes in excitability were quantified by measuring rheobase (minimum current required to elicit an action potential), determined by applying depolarizing current steps in 10 pA increments. The recording chamber was continuously perfused with an external solution at 2 ml/min. Solutions had the following composition (mM): pipette solution: K-gluconate 110, KCl 30, HEPES 10, MgCl_2_ 1, CaCl_2_ 2 (pH 7.2, adjusted with KOH; 290 mOsm); external solution: NaCl 143, KCl 5, HEPES 10, glucose 10, MgCl_2_ 1, CaCl_2_ 2 (pH 7.4, adjusted with NaOH; 305 mOsm). Neurons were pre-incubated with bacterial supernatant, diluted to a final concentration of 0.5X in external solution, for 10 min at RT before measuring rheobase. The following bacterial samples were tested: Mega Medium (10-fold concentrated, control), WT *B. fragilis* supernatant (10-fold concentrated), *B. fragilis* *∆bfp1* supernatant (10-fold concentrated). To determine the requirement for protease activity, supernatants were preincubated with the serine protease inhibitor FP-alkyne (100 µM, 60 min, RT) or vehicle before incubation with neurons. To determine the contribution of PAR_2_, neurons were pre-incubated with the PAR_2_ antagonist AZ3451 (1 µM, 60 min, RT) or vehicle before exposure to supernatant, or neurons from *Par_2_^-/-^* global KO mice were studied.

**Extracellular afferent nerve recording.** The distal colon of C57BL/6J wild-type mice, along with the intact neurovascular bundle of the inferior mesenteric artery, was removed and placed in a Sylgard-lined organ bath that was continuously superfused (10 mL/min) with oxygenated (5 % CO_2_-95 % O_2_) Krebs buffer solution (mM): 118.4 NaCl, 24.9 NaHCO_3_, 1.2 MgSO_4_, 1.2 KH_2_PO_4_, 11.7 glucose, and 1.9 CaCl_2_, pH 7.4 at 34 °C. The Krebs solution also contained the l-type calcium channel blocker nifedipine (3 µM), the muscarinic acetylcholine receptor antagonist atropine (5 µM), and the cyclooxygenase inhibitor indomethacin (3 µM). The colonic tissue preparation was then cannulated at both ends. The proximal end was attached to an infusion pump to allow continuous perfusion of the Krebs buffer solution (0.2 mL/min), while the distal end was linked to a pressure transducer that measured intracolonic pressure. The lumbar splanchnic nerve bundles emanating from the colon were separated into small strands and then drawn into a glass suction recording electrode attached to a Neurolog headstage (NL100; Digitimer). The signal was filtered through a 10 Hz low-cut filter to remove low-frequency noise and a 5000 Hz high-cut filter to remove high-frequency noise. The analogue signal was digitized with a Micro 1401 MKII interface and recorded using CED Spike2 6.1 software. A ramp distension was applied by closing the outflow drain of the preparation until the intracolonic pressure reached 60 mmHg to ensure the selected nerve bundle was mechanically sensitive. Nerve strands that did not display an increase in action potential (AP) frequency in response to distension were discarded. The mechanical distensions were performed every 15 min until the magnitude of the distension response was reproducible (defined as ≤ 20% change in maximal AP discharge frequency between three distensions). Following confirmation of nerve stability, Mega Medium or WT *B. fragilis* or *B. fragilis* *Δbfp1* supernatant was applied intraluminally by mixing 20μL of the 40x supernatant with 780 μL Krebs solution and perfusing for 10 min. At the end of the 10 min, but with supernatant still present in the colon, the afferent nerve response to ramp distension was reassessed. The intraluminal solution was then switched back to Krebs, and the colon was washed out for 30 min, distending every 15 min.  Single-unit analysis was used to discriminate individual afferent fibers.^10^ The analysis was conducted using the Spike2 sorting function. The basal firing rate was defined as the average spontaneous AP firing frequency (Hz) of individual axons (discriminated nerve units) in the absence of distensions. The basal AP discharge rate was measured for 120 s before the control distension and for 120 s just before the distension in the presence of supernatant.  The distension response immediately before the supernatant was used as the control distension. Nerve excitation in response to distensions performed in the presence of supernatant was compared with the control distension response to determine its impact on the magnitude and pressure-sensitivity of distension responses.

**Intracolonic administration of bacterial supernatants.** Investigators were blinded to treatments and genotypes. Mice were lightly sedated (3-5% isoflurane). The following samples were administered into the colon lumen by enema (150 µl, 3 cm from anus): vehicle (sterile sodium chloride 0.9%); Mega Medium (10-fold concentrated); WT *B. fragilis* supernatant (10-fold concentrated); or *B. fragilis* ∆*bfp1* supernatant (10-fold concentrated).

**Colonic pain.** Mice were acclimatized to the room, apparatus, and investigator for 2 h per day for 2 days before the study. The abdomen was divided into 9 equal quadrants. von Frey filaments of increasing force were applied to the central quadrant, corresponding to the colonic region. Responses to von Frey filament stimulation included arching of the back, jumping, and raising the rear legs. Responses were measured hourly for 6 h after intracolonic administration of samples. Results are expressed as a threshold in grams. To determine the contribution of PAR_2_ to pain, bacterial supernatants were similarly administered to *Par_2_^-/-^* global KO or *Par_2_^-/-^-Nav1.8* mice.

**Colonic inflammation.** The colon was removed 3 h after intracolonic administration of samples and was snap-frozen. RNA was extracted using a Direct-zol RNA Kit (Zymo Research, #R2050). cDNA was synthesized from 50 ng of DNase-treated RNA using a High-Capacity cDNA Reverse Transcriptase Kit (Applied Biosystems, #4374966). The cDNA (50 or 100 ng) was subjected to 40 cycles of qRT-PCR amplification using the QuantStudio 3 Real-Time PCR System. The mRNA expression of TNFα (#Mm00443258_m1), IL-1β (#Mm00434228_m1) and CXCL1 (#Mm04207460_m1) was measured by qPCR using TaqMan® gene expression and TaqMan® Fast Advanced Master Mix (#4444557). The relative amounts of target genes were calculated by normalizing the expression with the reference gene GAPDH (#Mm9999915_g1). The 2-ΔΔCT method was used to compare the reference and target gene levels.

**PAR_2_ endocytosis.** Samples were administered into the colon of *Par_2_-mugfp* mice as described above. The colon was removed and fixed (4% paraformaldehyde, PFA in PBS, 2 h, 4°C), cryoprotected (30% sucrose, PBS, 48 h, 4°C), and embedded in tissue freezing medium (TFM, #TFM-5, General Data). Frozen sections (8 µm) were prepared. Sections were blocked in 10% normal donkey serum (NDS), 0.05% Triton X-100 in PBS (1 h, RT). Sections were incubated with rabbit anti-GFP (1:400, overnight, 4°C; #600-401-215L, Rockland Immunochemicals). Slides were washed and incubated with donkey anti-rabbit Alexa Fluor^®^ 488 (1:1000, 45 min, RT; Invitrogen). Slides were incubated with DAPI (1 µg/ml, 5 min) and mounted in ProLong^®^ Gold Antifade (ThermoFisher). Sections were observed using a Leica SP8 confocal microscope with HCX PL APO 40x (NA 1.30) or 63x (NA 1.40) oil objectives (Leica-Microsystems). Images were processed using Adobe Photoshop and Illustrator.

**Antibiotic treatment and repopulation.** To deplete intestinal bacteria, mice received a mixture of antibiotics in drinking water / 10% sucrose for 7 d (days 0-7): ampicillin (1 g/L), vancomycin (0.5 g/L), neomycin (1 g/L) and metronidazole (1 g/L). During the repopulation phase (days 8-18), mice received vehicle (control), WT *B. fragilis* or *B. fragilis* *∆bfp1* (1x10^8^ cells per 100 μL) every other day by gavage (100 µL). The engraftment phase continued from day 18. Fecal pellets were collected every other day and were snap-frozen for assessing the depletion of gut bacteria and repopulation with *B. fragilis* by qRT-PCR. Fecal pellet bacterial DNA was extracted using the QIAmp® Power Fecal® Pro DNA Kit, (Qiagen, USA). Bacterial DNA was quantified by qRT-PCR using 16S primers (100 nmol/L) for total bacterial DNA (Universal_forward: AAACTCAAAKGAATTGACGG, Universal_reverse: CTCACRRCACGAGCTGAC) or for *B. fragilis* DNA (Bf_DNA_fwd: TGATTCCGCATGGTTTCATT, Bf_DNA_rev: CGACCCATAGAGCCTTCATC) with 2x Fast SYBR Green master mix (#43-856-12, Applied Biosystem, USA) and 15 ng bacterial DNA. qRT-PCR conditions were: denaturation for 10 min, 94°; 40 cycles of 1 min, 94°C; annealing for 1 min, 60°C; and elongation for 90 min, 72°C. *B. fragilis* DNA extracted from culture was used to plot a standard curve for both *B. fragilis* and universal primer sets, which was used to calculate bacterial DNA concentration. Abdominal withdrawal responses to stimulation with von Frey filaments were measured on day 0 before antibiotic treatment, day 7 after antibiotic treatment, and alternate days during the repopulation and engraftment phases.

**Spontaneous non-evoked behavior.** On day 19 of the engraftment, non-evoked behavior was assessed using the behavioral spectrometer (Behavior Sequencer, Behavioral Instruments). Mice were individually placed in the center of the behavioral spectrometer and their ambulatory, exploratory and grooming behaviors were recorded, tracked, evaluated, and analyzed using a computerized video tracking system (Viewer3, BiObserve) for 30 min. Total distance traveled in the open field, average velocity of locomotion, visit to the center of the arena, ambulation, and grooming were recorded and analyzed.

**Statistics.** Data are presented as mean ± SEM. For studies of cells, n>5 experiments were made. For experiments with mice, 5-6 mice were studied per treatment. Differences were assessed using Student's t-test for two comparisons and 1 or 2 way-ANOVA followed by Tukey for multiple comparisons test. p<0.05 was considered significant at the 95% confidence level.

**Supplementary Notes**

**Supplementary Note 1. Synthesis of Cys-PEG_4_-Cy5 and FP-probes**

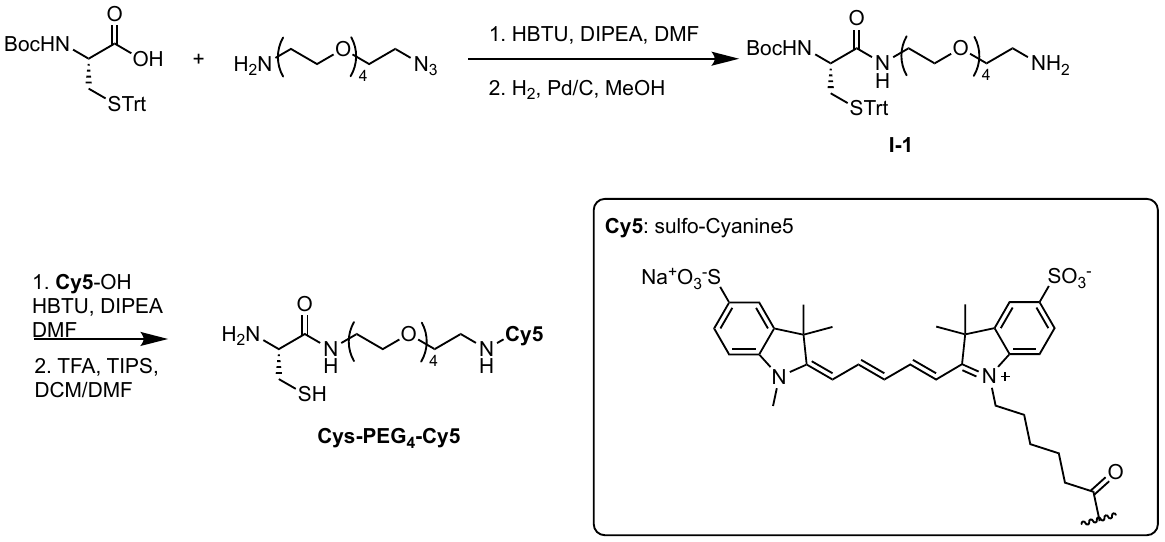

**Synthesis of I-1:** *N*-(tert butoxycarbonyl)-S-trityl-L-cysteine (160 mg, 0.35 mmol, 1.0 eq) and HBTU (144.0 mg, 0.38 mmol, 1.1 eq) were dissolved in DMF (3 mL). (2-[2-[2-[2-(2-Azidoethoxy)ethoxy]ethoxy]ethoxy]ethanamine (99.6 mg, 0.38 mmol, 1.1 eq) and DIPEA (89.2 mg, 0.69 mmol, 2.0 eq) were added and the solution was stirred at rt for 15 h. The solvent was removed under reduced pressure and the residue was redissolved in EtOAc. The organic layer was washed with 1 M citric acid, 5% NaHCO_3_ and brine and the solvent was removed under reduced pressure. The residue was dissolved in MeOH (4mL) and Pd/C (10%, 30 mg, 0.28 mmol) was added. Reduction was performed for 17 h using a hydrogen-filled balloon. The MeOH was removed under reduced pressure, the residue resuspended in EtOAc, filtered over celite and the solvent was removed under reduced pressure. Purification by preparative HPLC (A: H_2_O + 0.1% TFA, B: MeCN + 0.1% TFA, gradient: 5 to 95% B in 17 min) and lyophilization yielded the product as a white solid (205 mg, 0.26 mmol, 74%).

**Synthesis of Cys-PEG_4_-Cy5:** Sulfo-Cyanine5 (“Cy5”, 15 mg, 23 µmol, 1.0 eq) and HBTU (10 mg, 27 µmol, 1.2 eq) were dissolved in DMF (1 mL) and I-1 (18 mg, 23 µmol, 1.0 eq) and DIPEA (8.7 mg, 68 µmol, 3 eq) were added. The reaction was stirred overnight. Next, DCM (0.8 mL), TIPS (12 µL) and TFA (0.2 mL) were added and the reaction was stirred at rt for 60 min. The solvents were removed under reduced pressure and the residue was purified by preparative HPLC (A: H_2_O + 0.1% TFA, B: MeCN + 0.1% TFA, gradient: 5 to 95% B in 17 min), yielding the product as a blue powder after lyophilization (10 mg, 4.8 µmol, 20%).

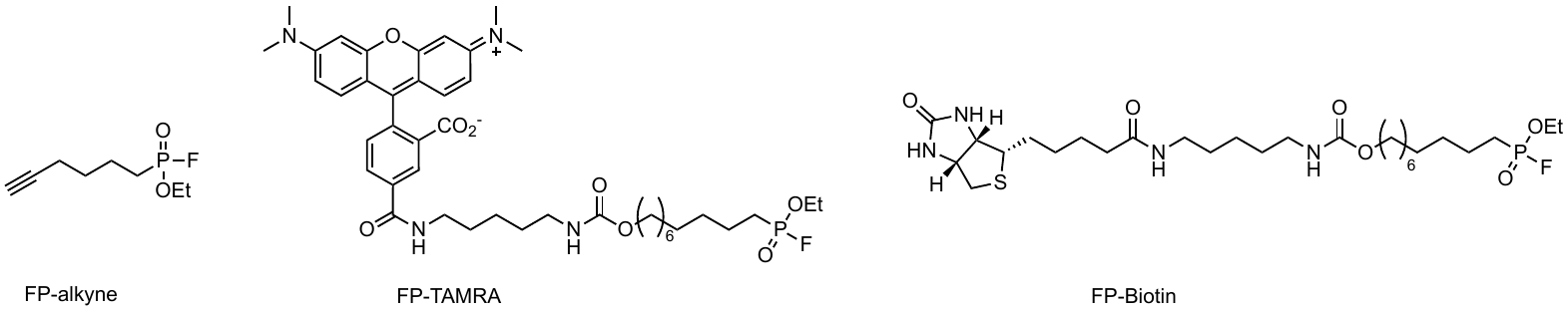

FP-alkyne was synthesized as previously reported.^11^

FP-TAMRA and FP-Biotin were synthesized as previously reported.^12^

**Supplementary Note 2. Preparation of Mega Medium**

The preparation of Mega Medium was performed as previously published:^1^

1. Add dry ingredients, liquid ingredients, and mix well (Table 1).
2. pH to ~7.0. In practice, this takes slightly less than 2.5 mL 10 M KOH per 500 mL media. After pH measurement, add 3.6 – X mL water where X is mL of KOH added.
3. If plates are needed, add 7.5 g agar per 500 mL of media.
4. Autoclave (121°C for 25 minutes at 20 PSI, 20 minutes dry).
5. As soon as autoclaving starts, remove vitamin mix from the freezer to thaw. Once thawed, make vitamin solution and filter sterilize (Table 2).
6. After media has cooled, sterilely add vitamin solution.

**Table 1.**

| **Component** | **Amount**  **(in 500 mL)** | **Final conc.** | **Mfr.** | **Vendor (cat. #)**  **[if any]** |
| --- | --- | --- | --- | --- |
| Milli-Q water (dH_2_O) | 410 mL |  |  |  |
| 1 M Potassium phosphate  buffer, pH 7.2^†^ | 50 mL | 10%  (v/v) |  |  |
| TYG salts solution^†^ | 20 mL | 4%  (v/v) |  |  |
| Tween 80 | 1 mL of 25% (v/v) | 0.05%  (v/v) |  |  |
| SCFA supplement^†^ | 1.4 mL | 0.28%  (v/v) |  |  |
| 0.8% (w/v) CaCl_2_^†^ | 500 µL |  |  |  |
| FeSO_4_·7H_2_O^†^ | 500 µL of 0.4 mg/mL |  |  |  |
| Resazurin^†^ | 2 mL of 0.25 mg/mL | 0.000001%  (w/v) | Sigma | Sigma (R2127) |
| Trypticase^TM^ Peptone | 5 g | 1%  (w/v) | BBL | BD (211921) |
| Yeast Extract | 2.5 g | 0.5%  (w/v) | Bacto | BD (212750) |
| Meat extract | 2.5 g | 0.5%  (w/v) | Sigma | Sigma (70164) |
| L-Cysteine hydrochloride | 0.25 g | 0.05%  (w/v) | Sigma | Sigma (C1276) |
| D-(+)-Glucose | 1 g | 0.2%  (w/v) | Sigma | Sigma (G8270) |
| D-(+)-Cellobiose | 0.5 g | 0.1%  (w/v) | Sigma | Sigma (C7252) |
| D-(+)-Maltose monohydrate | 0.5 g | 0.1%  (w/v) | Sigma | Sigma (M5885) |
| D-(-)-Fructose | 0.5 g | 0.1%  (w/v) | Sigma | Sigma (F0127) |

**Table 2.**

| Vitamin K solution^†^ | 500 µL of 1 mg/mL | 0.000001% (w/v) | Sigma | Sigma (M5625) |
| --- | --- | --- | --- | --- |
| Trace Mineral Supplement | 5 mL | 1% (v/v) | ATCC | ATCC (MD-TMS) |
| Vitamin Supplement | 5 mL | 1% (v/v) | ATCC | ATCC (MD-VS) |
| Histidine-Hematin^†^ | 500 µL |  |  |  |

^†^ STOCK SOLUTION RECIPES

**1 M potassium phosphate buffer, pH 7.2**

1. Prepare 1 M KH_2_PO_4_ (monobasic).

68.045 g KH_2_PO_4_ (anhydrous, f.w. = 136.09) in Milli-Q water to 500 mL

2. Prepare 1 M K_2_HPO_4_ (dibasic).

174.18 g K_2_HPO_4_ (anhydrous, f.w. = 174.18) in Milli-Q water to 1 L

3. Add monobasic to dibasic to achieve pH 7.2.

(You typically need ~430 mL monobasic added to 1 L dibasic)

*Tip: When preparing this solution, add the potassium phosphate powders to an actively stirring volume of water. Attempting to add water to a large mass of powder will result in the formation of a difficult to dissolve clump of material at the bottom of the bottle.*

**Vitamin K solution**

Dissolve 40 mg menadione (Vitamin K_3_, Sigma M5625) in 40 mL 100% EtOH.

**TYG salts solution**

MgSO_4_·7H_2_O (Sigma 230391)	0.5 g

NaHCO_3_ (Sigma S5761)	10.0 g

NaCl (Sigma S7653)	2.0 g

Milli-Q water	to 1 L

**FeSO_4_·7H_2_O (0.4 mg/mL)**

Dissolve 40 mg FeSO_4_·7H_2_O (Sigma F8633) in 100 mL Milli-Q water.

**0.8% (w/v) CaCl_2_**

Dissolve 0.4 g CaCl_2_·2H_2_O (Sigma C7902) in 50 mL Milli-Q water.

**Resazurin anaerobic indicator (0.25 mg/mL)**

1. Dissolve 25 mg resazurin (Sigma R2127) in 100 mL distilled H_2_O.
2. Store protected from light at 4 °C.

**Histidine-Hematin**

1. Prepare 0.2 M histidine, pH 8.0

- Mix 4.2 g Histidine-HCl monohydrate (Sigma H7875) in 80 mL Milli-Q water.
- Adjust the pH from 4 to 8 with 10 N NaOH (the histidine will go into solution as the pH rises).
- Bring the final volume to 100 mL with Milli-Q water.

2. Mix 12 mg hematin (Sigma H3281) with 10 mL of 0.2 M histidine, pH 8.0. Dissolve by end-over-end rotation or vigorous shaking for several hours. Filter-sterilize using 0.2 µm filter.

**SCFA supplement**

Acetic acid, glacial (Sigma A6283)	17 mL

Propionic acid (Sigma P5561)	6 mL

Butyric acid (Sigma B103500)	4 mL

Isovaleric acid (Sigma 129542)	1 mL

**Supplementary References**

1. Han, S., Treuren, W.V., Fischer, C.R., Merrill, B.D., DeFelice, B.C., Sanchez, J.M., Higginbottom, S.K., Guthrie, L., Fall, L.A., Dodd, D., et al. (2021). A metabolomics pipeline for the mechanistic interrogation of the gut microbiome. Nature *595*, 415–420. <https://doi.org/10.1038/s41586-021-03707-9>.

2. Cox, J., and Mann, M. (2008). MaxQuant enables high peptide identification rates, individualized p.p.b.-range mass accuracies and proteome-wide protein quantification. Nat. Biotechnol. *26*, 1367–1372. <https://doi.org/10.1038/nbt.1511>.

3. Tyanova, S., Temu, T., Sinitcyn, P., Carlson, A., Hein, M.Y., Geiger, T., Mann, M., and Cox, J. (2016). The Perseus computational platform for comprehensive analysis of (prote)omics data. Nat. Methods *13*, 731–740. <https://doi.org/10.1038/nmeth.3901>.

4. Zou, Y., Xue, W., Luo, G., Deng, Z., Qin, P., Guo, R., Sun, H., Xia, Y., Liang, S., Dai, Y., et al. (2019). 1,520 reference genomes from cultivated human gut bacteria enable functional microbiome analyses. Nat. Biotechnol. *37*, 179–185. <https://doi.org/10.1038/s41587-018-0008-8>.

5. Camacho, C., Coulouris, G., Avagyan, V., Ma, N., Papadopoulos, J., Bealer, K., and Madden, T.L. (2009). BLAST+: architecture and applications. BMC Bioinform. *10*, 421. <https://doi.org/10.1186/1471-2105-10-421>.

6. García-Bayona, L., and Comstock, L.E. (2019). Streamlined Genetic Manipulation of Diverse Bacteroides and Parabacteroides Isolates from the Human Gut Microbiota. Mbio *10*, e01762-19. <https://doi.org/10.1128/mbio.01762-19>.

7. Co, J.Y., Margalef-Català, M., Li, X., Mah, A.T., Kuo, C.J., Monack, D.M., and Amieva, M.R. (2019). Controlling Epithelial Polarity: A Human Enteroid Model for Host-Pathogen Interactions. Cell Reports *26*, 2509-2520.e4. <https://doi.org/10.1016/j.celrep.2019.01.108>.

8. StemCellTechnologies How to Generate Human Intestinal Organoid-Derived Monolayers Using IntestiCult. <https://www.stemcell.com/how-to-generate-human-intestinal-organoid-derived-monolayers-using-intesticult.html>.

9. Tu, N.H., Jensen, D.D., Anderson, B.M., Chen, E., Jimenez-Vargas, N.N., Scheff, N.N., Inoue, K., Tran, H.D., Dolan, J.C., Meek, T.A., et al. (2021). Legumain Induces Oral Cancer Pain by Biased Agonism of Protease-Activated Receptor-2. J. Neurosci. *41*, 193–210. <https://doi.org/10.1523/jneurosci.1211-20.2020>.

10. Nullens, S., Deiteren, A., Jiang, W., Keating, C., Ceuleers, H., Francque, S., Grundy, D., Man, J.G.D., and Winter, B.Y.D. (2016). In Vitro Recording of Mesenteric Afferent Nerve Activity in Mouse Jejunal and Colonic Segments. J. Vis. Exp. : JoVE. <https://doi.org/10.3791/54576>.

11. Keller, L.J., Nguyen, T.H., Liu, L.J., Hurysz, B.M., Lakemeyer, M., Guerra, M., Gelsinger, D.J., Chanin, R., Ngo, N., Lum, K.M., et al. (2023). Chemoproteomic identification of a DPP4 homolog in Bacteroides thetaiotaomicron. Nat. Chem. Biol. *19*, 1469–1479. <https://doi.org/10.1038/s41589-023-01357-8>.

12. Liu, Y., Patricelli, M.P., and Cravatt, B.F. (1999). Activity-based protein profiling: The serine hydrolases. Proc. Natl. Acad. Sci. *96*, 14694–14699. <https://doi.org/10.1073/pnas.96.26.14694>.
